# Beyond Co-Existence: Early *α*-Synuclein-Amyloid-*β*_42_ Association from Molecular Dynamics Simulations

**DOI:** 10.64898/2026.01.29.702600

**Authors:** F. Carvalho, P. Maximiano, M. Hashemi, P. N. Simões

**Affiliations:** University of Coimbra, CERES, Department of Chemical Engineering, Rua Sílvio de Lima, Coimbra, 3030-790, Portugal; Department of Physics, Auburn University, Leach Science Center 3126, Auburn, AL, 36849-5319, USA

**Keywords:** Protein aggregation, Alpha-synuclein, Amyloid-beta, Molecular Dynamics, Hybrid oligomers

## Abstract

Amyloid-*β* (A*β*) deposits often appear in Parkinson’s disease and *α*-synuclein (*α*-syn) pathology in Alzheimer’s disease. As soluble oligomers contribute importantly to cytotoxicity, their early crosstalk is critical to understand. Using all-atom molecular dynamics, we investigated how the initial oligomeric configuration of A*β*_42_ shapes its early association with *α*-syn, simulating *α*-syn with (i) one A*β*_42_ monomer, (ii) two separated monomers, and (iii) a pre-formed dimer plus a free monomer, across 15 independent 3 µs trajectories. Association patterns depended strongly on the initial configuration. With monomeric or nascent dimeric A*β*_42_, *α*-syn frequently contacted the metal-binding, central hydrophobic core (CHC), and C-terminal regions, and these contacts were temporally coordinated, most reproducibly the *α*-syn NAC with the A*β*_42_ CHC and polar regions. *β*-hairpin sampling was reduced specifically when *α*-syn engaged the CHC and C-terminus, an interface-dependent rather than a general effect that did not track contact duration, and coincided with more expanded conformations. With a pre-formed dimer, *α*-syn associated mainly peripherally, while the dimer’s *β*-structured interface remained comparatively stable. First-order dihedral entropy differed only modestly between the free and associated states, indicating that association redistributed conformational populations without globally reducing local backbone diversity. Overall, *α*-syn’s early structural effect on A*β*_42_ is configuration-dependent: it engages and perturbs lower-order species while associating only peripherally with the sampled dimer.

## 1. Introduction

Alzheimer’s disease (AD) and Parkinson’s disease (PD) are traditionally associated with distinct protein aggregates: amyloid-*β* (A*β*) plaques and tau neurofibrillary tangles in AD, and *α*-synuclein (*α*-syn)-rich Lewy bodies in PD [1–4]. However, this pathological separation is increasingly being challenged by clinical and neuropathological evidence. A substantial fraction of AD cases exhibits *α*-syn pathology, whereas A*β* deposits are also observed in PD and related synucleinopathies [3, 5]. This comorbid pathology is associated with more severe clinical outcomes, including accelerated cognitive decline and increased mortality, compared to “pure AD” [4, 6]. Importantly, A*β* and *α*-syn appear to interact directly rather than simply accumulate in parallel, suggesting that their molecular crosstalk may contribute to disease progression through conformation-dependent mechanisms [3, 7–9].

Among A*β* isoforms, A*β*_42_ is particularly relevant because it is more aggregation-prone and neurotoxic than A*β*_40_, which is the most abundant [10, 11]. A*β*_42_ is intrinsically disordered and samples a heterogeneous conformational ensemble, but early aggregation is thought to depend on rare monomeric states that promote stable nucleus formation [5, 12, 13]. The functional A*β*_42_ regions are defined as metal-binding (residues 1–16), CHC (central hydrophobic core, residues 17–21), polar (residues 22– 29), and C-terminus, residues 30–42 [14]. Transient contacts between the CHC and hydrophobic C-terminus can favor *β*-hairpin-like conformations, in which two *β*-strand segments are connected by a short turn [15–17]. These motifs are considered important early structural elements in A*β*_42_ nucleation and oligomer growth. Because soluble oligomeric species are strongly implicated in synaptic dysfunction and cytotoxicity, understanding how these early conformations are stabilized, disrupted, or redirected is central to understanding A*β*_42_ toxicity [10, 11].

*α*-syn is a 140-residue presynaptic intrinsically disordered protein (IDP) composed of three major regions: an amphipathic N-terminal domain (residues 1–60), a hydrophobic non-amyloid component (NAC) domain (residues 61–95), and an acidic C-terminal tail (residues 96–140) [18–20]. These domains provide different possible interaction modes with A*β*_42_. Electrostatic contacts may occur between the lysine-rich N-terminus of *α*-syn and acidic residues in A*β*_42_, whereas hydrophobic interactions may involve the *α*-syn NAC and CHC/C-terminal regions of A*β*_42_ [21, 22]. The NAC region was originally identified in A*β* plaques, and later biochemical, biophysical, and computational studies support the possibility of a direct physical association between these two aggregation-prone polypeptides [3, 18, 23].

The functional consequences of A*β*_42_–*α*-syn association remain heterogeneous. Several studies suggest that *α*-syn can sequester A*β*_42_ monomers, inhibit mature fibril formation, and redirect A*β*_42_ toward soluble hetero-oligomeric assemblies [24–28]. Conversely, A*β*_42_ can alter *α*-syn conformation and promote *α*-syn aggregation [3, 25]. In this context, shielding and scaffolding are most usefully treated as conceptually limiting behaviors rather than as mutually exclusive or directly predictive mechanisms. A shielding-like interaction denotes the occupation by *α*-syn of A*β*_42_ surfaces that could otherwise contribute to intramolecular or intermolecular amyloidogenic contacts. A scaffold-like interaction denotes an association with an existing or developing A*β*_42_ assembly without immediate disruption of its principal interface. An individual encounter complex may display features of either behavior or both, depending on the initial A*β*_42_ configuration, the regions engaged, and the geometry of the interface [29–32]. Therefore, the present simulations were designed to identify interaction patterns compatible with these limiting behaviors, not to establish a universal transition criterion between them.

Molecular dynamics (MD) simulations have become indispensable for examining these transient and heterogeneous encounter complexes at an atomistic resolution [13, 33]. Previous computational studies have characterized A*β*_42_–*α*-syn heterodimers and identified possible electrostatic and hydrophobic interfaces[21, 22]. Atomistic simulations have also helped clarify how specific *α*-syn domains, particularly the N-terminal and NAC regions, contribute to the early A*β*_42_–*α*-syn association [34–36]. However, little is known about how *α*-syn interacts with A*β*_42_ once A*β*_42_ begins to self-associate. This distinction is important mechanistically. Binding to two initially separated A*β*_42_ monomers probes how *α*-syn associates during nascent dimer formation, whereas binding to a pre-formed A*β*_42_ dimer probes whether *α*-syn contacts peripheral or core-facing surfaces and whether detectable remodeling occurs within the simulated time scale.

Here, we use all-atom MD simulations to examine the early dynamics of *α*-syn interacting with three A*β*_42_ configurations: one A*β*_42_ monomer, two initially separated A*β*_42_ monomers, and a pre-formed A*β*_42_ dimer plus a free monomer. Using five independent 3 µs replicates per system, we characterized the encounter complexes and immediate conformational changes accessible within the simulated timescale. We quantified aggregation events, binding hierarchies, contact interfaces, *β*-hairpin formation, and contact-dependent structural changes. A first-order Shannon entropy analysis of the backbone dihedral distributions was used as a descriptive measure of local conformational diversity. The comparison is deliberately configuration-specific; it tests how the three selected starting arrangements differ rather than representing the full ensemble of A*β*_42_ oligomers, stoichiometries, or physiological concentrations.

## 2. Methods

### System Setup and Simulation Parameters

Three model systems were constructed to investigate the aggregation hierarchy: System 1, containing one *α*-syn and one A*β*_42_ monomer 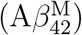; System 2, containing one *α*-syn and two A*β*_42_ monomers (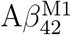 and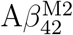); and System 3, containing one *α*-syn, one A*β*_42_ monomer 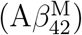, and one pre-formed A*β*_42_ dimer (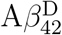, with individual A*β*_42_ chains denoted as 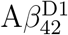 and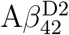).

The initial structures for *α*-syn and 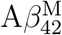 were generated to mimic the experimental constructs that are often used in single-molecule studies. Regarding the initial structure of the 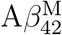 chain, an additional cysteine (CYS) residue was included in the metal-binding region (position 0) to mimic experimental conditions, where the presence of CYS is used to attach the peptide to substrates or fluorophores for AFM or fluorescence studies, respectively. Controls have shown that the additional cysteine does not affect the 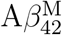 behavior or aggregation process[4]. The coordinates for Cys-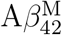 were obtained from simulation studies of amyloid-cholesterol interactions[37]. To evaluate how *α*-syn interacts with structured amyloid aggregates, 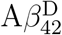 (System 3) was extracted from a previous self-assembly study[38]. The pre-formed A*β*_42_ dimer represents a structurally specific starting configuration selected to contrast an established A*β*_42_–A*β*_42_ interface with initially separated monomers. It is not intended to represent the equilibrium distribution of A*β*_42_ dimers. Conclusions involving core accessibility, peripheral binding, and resistance to remodeling apply only to the sampled geometry and serve as a case study of oligomer-state-dependent association.

The starting structure for *α*-syn was adopted from the solution ensemble described by Robustelli et al.[39]. All systems were solvated in a rectangular box with dimensions 17 *×* 17 *×* 14 nm using the TIP3P water model[40] (approximately 126 000 water molecules per system). To neutralize the system charges and mimic physiological conditions, Na^+^ and Cl^*−*^ were added at a concentration of 150 mM.

Simulations were performed using GROMACS[41] with the CHARMM36m force field[42]. The protocol consisted of energy minimization (steepest descent, *<*1000 kJ mol^−1^ nm^−1^) followed by 1 ns NVT and 1 ns NPT equilibration runs. During equilibration, position restraints were applied to the protein backbone (*k* = 400 kJ mol^−1^ nm^−2^) and side chains (*k* = 40 kJ mol^−1^ nm^−2^). Production runs were performed for 3 *µ*s per replicate. All polypeptide species were placed in random relative positions for each of the five replicates of the three systems, while ensuring a minimum distance of 4 nm between adjacent polypeptides. The temperature was maintained at 300 K using the V-rescale thermostat[43] (*τ*_*t*_ = 1.0 ps), and pressure was coupled isotropically at 1.0 bar using the Parrinello-Rahman barostat[44] (*τ*_*p*_ = 5.0 ps, compressibility 4.5 × 10^−5^ bar^−1^). Long-range electrostatics were treated using PME[45], and short-range nonbonded interactions were cut off at 1.2 nm with a force switch starting at 1.0 nm. Hydrogen bonds were constrained using LINCS[46] to allow an integration step of 2 fs.

### Trajectory Processing

Prior to analysis, the solvent-stripped trajectories were processed using CPPTRAJ[47] to eliminate periodic boundary condition (PBC) artifacts and remove global translation and rotation. The processing workflow consists of two distinct stages.

First, PBC corrections were applied using the `autoimage` and `fiximagedbonds` commands to correct the split bonds, rewrap all molecules into the primary unit cell, and center the system. To ensure consistency across varying stoichiometries, the centering anchor was defined as the first A*β*_42_ chain in the topology (the N-terminal cysteine plus residues 1–42). This corresponds to the solitary 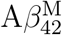 in Systems 1 and 3 and 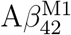 (distinguished from monomer 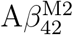) in System 2.

Second, to facilitate the visualization and analysis of relative inter-protein motions, frame-by-frame RMS alignment was performed (rms :1-43@CA previous). This aligned the C_*α*_ atoms of the anchor peptide in each frame to the preceding frame, effectively removing the rotational and translational drift while preserving the internal dynamics of the complex.

### Convergence Diagnostics

Full-trajectory convergence diagnostics were employed to evaluate the conformational sampling across the 3 µs trajectories. For each independent replicate, temporal evolution was assessed across the following structural metrics: intermolecular heavy-atom contacts, minimum inter-chain distance, and the radius of gyration for both *α*-syn and A*β*_42_ monomer chains. Sampling was characterized using three standard time-series analyses: (1) block averaging, dividing each trajectory into ten equal tem-poral blocks to estimate the local sampling variance and standard error; (2) cumulative running averages to confirm the stabilization of the global ensemble mean over the simulation timescale; and (3) temporal autocorrelation functions to evaluate the timescale of conformational memory and verify the sampling of statistically independent states. The resulting composite diagnostics for all systems and replicates are provided in the Supporting Information (Section I).

### Aggregation Events

An interaction was defined as an aggregation event only if the frame simultaneously satisfied a minimum distance of ≤ 0.25 nm between any atoms of the interacting species and a total of ≥ 50 intermolecular heavy atoms contacts (defined as any pair of heavy atoms within 0.6 nm). This dual-criteria metric was defined by plotting heavy atom contacts versus the minimum distance (results not shown) and was employed to eliminate the false positives inherent to the single-parameter definitions. A strict minimum-distance criterion alone misclassifies transient single-atom glancing blows as aggregates, whereas a pure contact-count criterion (*≤*0.6 nm) can capture loose complexes separated by a hydration layer. This simultaneous requirement guarantees that the polypeptides physically dock and establish a substantial interaction interface.

### Intermolecular Contact Maps

To map the binding interfaces, residue-wise contact matrices were calculated using CPPTRAJ. An active contact was strictly defined between any heavy atom of residues *i* and *j* within a distance cutoff of 0.6 nm. For every residue pair across the interacting polypeptides, the fractional contact time was calculated by dividing the number of frames in which at least one active contact existed by the total number of evaluated simulation frames (30 000).

To quantify domain-level interactions, these residue-level fractional contact times were summed over established polypeptide structural regions based on known functional motifs[14, 20]. For A*β*_42_, residues are reported using their canonical A*β*_42_ numbering. The N-terminal cysteine tag is assigned position 0 and is grouped into the metal-binding domain, so that all native A*β*_42_ residues retain their canonical numbers.

To account for the inter-replicate heterogeneity inherent in the association of intrinsically disordered proteins, ensemble-averaged contact frequencies were accompanied by 95% confidence intervals. These were calculated using a nonparametric bootstrap resampling approach: for each region-level pair, the five independent replicate matrices were resampled with replacement 1000 times to generate a distribution of the mean contact frequency. The lower and upper bounds of the 95% confidence interval were determined from the 2.5th and 97.5th percentiles of this distribution, respectively, with the lower bound constrained to a physical minimum of 0 %.

### Temporal Coordination Between Contact Regions

The contact matrices report each region-level pair as a trajectory average and cannot indicate whether two pairs engage concurrently. To resolve this, the number of active residue contacts within each pair of structural regions was recorded per frame using the previously described contact definition and regional assignments; the trajectory average of this count is the cumulative contact index reported previously. Frames with no active intermolecular contact were excluded, so that the analysis describes the composition of an existing interface rather than its formation.

Following correlation analysis of interatomic contacts in molecular dynamics [48, 49], the covariation of contact counts from two region-level pairs was quantified by the Pearson partial correlation coefficient *r*; values of +1, 0 and *−*1 denote that the pairs gain and lose contacts in proportion, vary independently, or vary oppositely. Ordinary correlation, as implemented in tools such as PyContact [50], cannot separate the genuine concerted engagement between single pairs from the collective variations in all pairs as the interface tightens and loosens. As such, the coefficient is made *partial*, isolating the covariation not accounted for by overall engagement [51, 52]. Engagement was measured from the region-level pairs other than the two compared, since a total containing them would force an artifactual negative correlation. As uniform tightening produces only positive coefficients, positive values are reported above *r* = 0.20, the level it reaches alone, and negative values at any magnitude.

As for the contact map, coefficients were computed per replicate and reported as the mean over the five replicates, with 95% confidence intervals from a nonparametric bootstrap over the five replicate coefficients (percentile method, 1000 resamples); frames were not pooled across replicates. Because the residues engaged in the A*β*_42_– *α*-syn association vary between replicates while the regions do not, all A*β*_42_ species were pooled, so the coefficients describe *α*-syn–A*β*_42_ region coordination irrespective of which A*β*_42_ chain is engaged, and establish which region pairs engage concurrently or alternatively rather than measuring interaction strength.

### Secondary Structure and β-Hairpin Analysis

The time-dependent secondary structure evolution was computed using the Define Secondary Structure of Proteins (DSSP) algorithm [53] implemented in CPPTRAJ. *β*-hairpin motifs within the A*β*_42_ peptide were initially identified using a search algorithm based on secondary structure assignments, defining a hairpin as two anti-parallel *β*-strands (DSSP residues E or B, length ≥ 2 residues) separated by a short turn or loop (length 2–5 residues). The strand length constraint ensures the presence of minimal cooperative hydrogen bonding required for a stable secondary structure [54], whereas the loop definition aligns with the geometries established by Sibanda and Thornton [55]. To eliminate false positives, these structural candidates were validated in 3D space, requiring a minimum of 15 heavy-atom contacts (distance *≤* 0.45 nm) between the two strands. Finally, to isolate amyloidogenic folding events relevant to aggregation, the analysis specifically filtered for *β*-hairpins, where both strands involved the core hydrophobic regions (CHC, residues 17–21, and/or the C-terminus, residues 30–42). This region-specific filtering is empirically justified by our simulations of the pre-formed A*β*_42_ dimer (System 3), which demonstrated that *>* 99% of persistent, mature *β*-hairpins are localized exclusively to these hydrophobic domains. To prevent double-counting of simultaneous structural registers, the presence of at least one validated region-specific motif was tracked on a per-frame basis over the simulation time to monitor the structural maturation of A*β*_42_.

### Structural Ensemble Analysis of Aβ_42_ and α-syn

To quantify the conformational consequences of inter-chain association, a structural ensemble analysis was performed on each A*β*_42_ and *α*-syn chain across all 15 trajectories. For each frame of each replicate, chains were classified into mutually exclusive contact states using the same dual-criteria metric applied in the aggregation event analysis (minimum inter-chain distance *≤*0.25 nm and ≥50 intermolecular heavy-atom contacts within 0.6 nm).

For each classified frame in each state, the structural metrics were computed per chain using MDAnalysis[56]. For A*β*_42_, these comprised the radius of gyration (*R*_g_), the *β*-strand fraction derived from DSSP assignments computed previously with CPPTRAJ, and the intra-C-terminal C*α*–C*α* distance between residues 30 and 42, which quantifies the degree of C-terminal compaction. For *α*-syn, *R*_g_ and *α*-helix fraction were computed alongside the C*α*–C*α* distance between the NAC region center (residue 78) and the C-terminus center (residue 118), which reports on the NAC–C-terminus proximity implicated in *α*-syn self-assembly. Distributions were compared using two-sided Mann-Whitney U tests applied to pooled frame ensembles. To control the family wise error rate and prevent false positives, *p*-values were corrected for multiple tests using the Bonferroni method across a pre-specified set of independent comparisons. Specifically, the correction was applied dynamically per metric: across 10 pre-specified comparisons for universal metrics (such as *R*_g_ and secondary structure fractions), 7 comparisons for A*β*_42_-specific spatial metrics (such as intra-C-terminal distance), and 3 comparisons for *α*-syn-specific spatial metrics (such as NAC–C-terminus distance). Effect sizes were quantified using the median difference and Cohen’s *d*.

### Time-resolved Configurational Entropy

To relate aggregation to the conformational freedom of each chain, we evaluated the first-order configurational entropy from the backbone’s dihedral angles. Following the mutual information expansion of Killian, Kravitz, and Gilson [57], the total configurational entropy of a chain can be approximated by the sum of single-coordinate entropies reduced by mutual information terms for correlations between coordinates. Here, we utilized its leading term, *S*_1_ = Σ_*i*_ *S*_*i*_, which represents the sum of the marginal entropies of the backbone (*ϕ, ψ*) dihedrals. Internal coordinates were selected because they are invariant to overall translation and rotation, requiring no structural superposition. This makes dihedral-based entropy measures particularly appropriate for intrinsically disordered chains such as *α*-syn and A*β*_42_ [58]. Each marginal entropy *S*_*i*_ was computed from the normalized histogram of its dihedral (36 bins over the 360° range) and scaled by the universal gas constant to report the values in kJ mol^−1^ K^−1^.

To resolve conformational freedom in time, *S*_1_ was evaluated locally in fixed-width sliding windows along each trajectory (20 ns; 200 frames; advanced in 5 ns steps), yielding a time series of the conformational diversity sampled by each chain. Because the bias of an entropy estimate depends on the number of effectively independent samples, scaling with the block length for a contiguous block [59], holding the window length strictly fixed ensures that the bias is identical for windows drawn from different aggregation states. This allows the bias to effectively cancel out when states are compared, permitting the windowed *S*_1_ to be interpreted accurately on a relative scale without requiring further finite sample corrections.

Each frame was assigned to a single aggregation state based on the contact definitions utilized in the aggregation analysis (heavy-atom minimum distance *≤* 0.25 nm with ≥ 50 heavy-atom contacts). These states were made mutually exclusive via a highest-order assembly priority (ternary *>* binary *>* free), and each window inherited the majority state of its constituent frames. For every chain, environment, and state, the median windowed *S*_1_ was computed for each replicate. The mean of these per-replicate medians *±* standard error across replicates is reported, ensuring that frames were not pooled across independent replicates. A lower windowed *S*_1_ in an aggregated state reflects reduced conformational diversity sampled per unit time, combining both a restricted accessible ensemble and slower exploration, which the present sampling does not uncouple. Consequently, this metric is interpreted descriptively in terms of structural flexibility rather than as a strict thermodynamic entropy difference.

### Visualization

Molecular snapshots and structural overlays were generated using Visual Molecular Dynamics (VMD)[60] and rendered using a Tachyon ray tracer.

## 3. Results and Discussion

The reliability of MD findings depends critically on the accuracy of the underlying force fields (FF), particularly for IDPs, which populate a flat and rugged energy landscape [14]. Standard FF often over-stabilize compact, globular structures and fail to capture the extended ensembles typical of IDPs. To address this, recent comparative studies have utilized modified FF, such as CHARMM36m and AMBER99SB-disp, which have been specifically reparameterized to balance water-protein interactions and prevent artificial collapse [33, 61]. Recent benchmarks comparing these FF have shown that while implicit solvent models may induce excessive compaction, explicit solvent simulations utilizing CHARMM36m successfully reproduce the disordered nature of monomeric A*β*_42_ and *α*-syn, matching experimental radii of gyration (*R*_g_) and NMR observables [22]. The simulations from this study, which are performed in an aqueous environment, will be compared with simulations of the exact same systems in an environment containing a lipid bilayer in a future work. Crucially, the CHARMM36m force field is fully compatible with the validated CHARMM36 lipid force field [62]. To ensure an accurate thermodynamic and conformational comparison of the role of the bilayer in heterotypic association, the selection of CHARMM36m for the water environment, whose results act as a control for the more complex lipid bilayer environment, was strictly motivated by these parallel bilayer simulations.

We quantified the interaction between *α*-syn and different species of A*β*_42_ to determine how the assembly state of A*β*_42_ influences its early association with *α*-syn. We analyzed three all-atom MD simulation systems: *α*-syn with one A*β*_42_ monomer, *α*-syn with two initially separated A*β*_42_ monomers, and *α*-syn with one A*β*_42_ monomer plus a pre-formed A*β*_42_ dimer. Each system was simulated using five independent 3 µs replicates. Convergence diagnostics, including block averaging, cumulative averages, and autocorrelation analysis, are provided for all systems and replicates in Supporting Information Section I. These diagnostics indicate that the simulations captured stochastic encounters and aggregation events within the accessible microsecond timescale, consistent with the exploratory scope of this study.

### 3.1 Aggregation events

Figure 1 summarizes the contact-defined aggregation events for the three systems. In every system, *α*-syn formed repeated contacts with A*β*_42_, and the evolution of these assemblies appeared to be governed by the initial oligomeric configuration of A*β*_42_.

**Figure 1.**
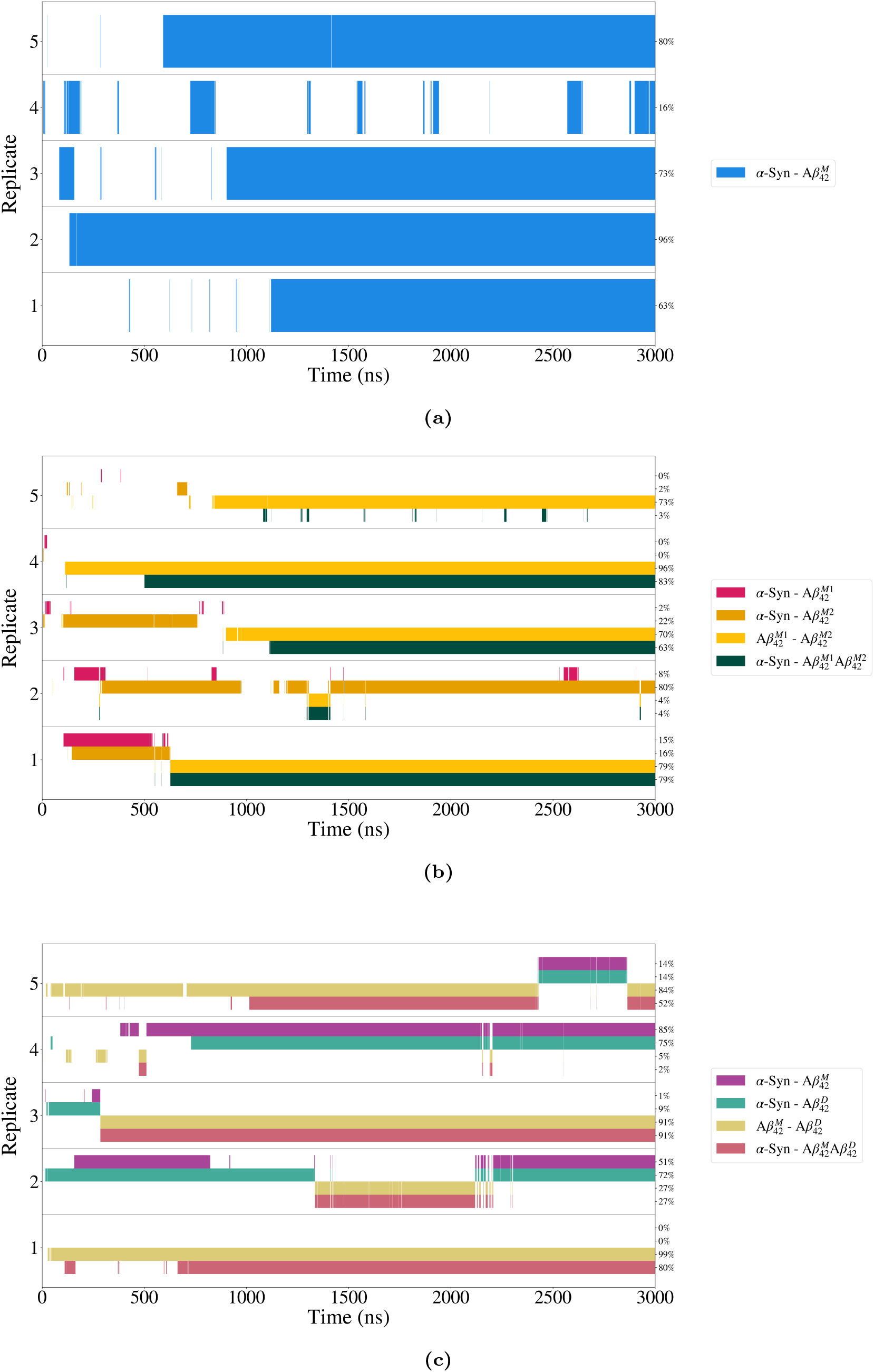
Aggregation event timelines. **(a)** System 1: 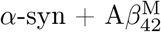. **(b)** System 2: 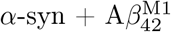 and 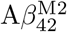. **(c)** System 3: 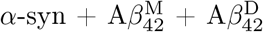. Each timeline shows aggregation events timelines across five replicates over 3 µs simulations.

In System 1, which contained one *α*-syn and one 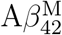 (Figure 1a), aggregation was observed in all five replicates. However, the persistence of the *α*-syn–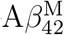complex varied substantially, ranging from 16% to 96% of the trajectory time. Replicates 2 and 3 formed long-lived complexes early in the simulation, with contact occupancies of 96% and 73%, respectively. In contrast, replicate 4 showed repeated association and dissociation events, with only 16% contact occupancy. This replicate-to-replicate variability suggests that the *α*-syn–A*β*_42_ heterodimer does not adopt a single dominant encounter geometry on this timescale. Instead, the interface samples multiple local association states, consistent with the conformational heterogeneity expected for two intrinsically disordered polypeptides and with previous binding free-energy calculations for A*β*_42_– *α*-syn complexes [22]. The broad range of complex persistence is also consistent with the entropic cost associated with the formation of stable interfaces between disordered monomers, as described for the rugged “funnel to disorder” landscape of monomeric A*β*_42_ [15].

The addition of a second A*β*_42_ monomer in System 2 introduced competition between A*β*_42_ homodimerization and heteromeric association with *α*-syn (Figure 1b). In

3 out of 5 replicates, 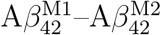 contacts occurred together with *α*-syn– 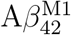or *α*-syn– 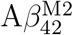 contacts, indicating that A*β*_42_ self-association and *α*-syn binding can occur within the same trajectory. In replicates 1, 3, and 4, a three-chain *α*-syn– 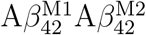 complex emerged and persisted for a large fraction of the simulation, suggesting that *α*-syn can remain associated with a nascent A*β*_42_ dimer once the two A*β*_42_ monomers form a stable interface. Replicates 2 and 5 showed different behaviors: in the former, the A*β*_42_ homodimer appeared only transiently, while *α*-syn remained primarily asso-ciated with 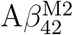, whereas in the later, 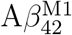 and 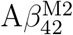 formed a stable homodimer early in the simulation, with 73% occupancy, whereas *α*-syn showed little association with this complex, with only 3% contact time. Replicate 5 represents an illustrative case in which A*β*_42_ self-association proceeded with minimal *α*-syn engagement. We return to this trajectory in Section 3.3, where it shows a continuous *β*-hairpin signature. Together, these trajectories highlight the kinetic heterogeneity of the three-chain system and suggest that *α*-syn engagement depends on the conformational accessibility of the forming A*β*_42_ dimer.

System 3 (Figure 1c) contained a pre-formed A*β*_42_ dimer and a free monomer. *α*-syn maintained stable contact with at least one A*β*_42_ species for at least 51% of each trajectory. In this system, *α*-syn– 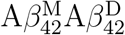 trimer formation occurs, but the lifetime of the trimer varies across replicates. In replicates 1, 3, and 5, the *α*-syn– 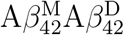 trimer complex existed at 80%, 91%, and 52%, respectively. The aggregation events observed in System 3 confirm that *α*-syn can actively bind to A*β*_42_ dimers or trimers (in addition to monomers) and engage a free monomer in parallel. These findings align with the oligomer stabilization mechanism described by Chau and Kim [24] and are of crucial relevance, as A*β*_42_ oligomers are established as the primary neurotoxic drivers, surpassing mature fibrils in terms of pathogenicity [13, 63]. Additionally, the formation of higher-order oligomers mirrors the broader literature on co-pathology. Early computational models by Tsigelny et al. [64] proposed that *α*-syn could dock onto membrane-bound A*β* oligomers, forming hybrid ring-like pentamers that exacerbate membrane permeabilization. Recently, Huang et al. [36] demonstrated that the co-aggregation of *α*-syn with A*β* stabilizes *β*-sheet-rich oligomers and facilitates the formation of toxic *β*-barrel intermediates, a structural motif often associated with pore formation and cytotoxicity in neurons. Our observation of stable hybrid trimers in System 3 provides atomistic support for this mechanism, which will be further explored in terms of the binding interface and *β*-hairpin content.

### 3.2 α-syn and Aβ_42_ contact Interface

Aggregation timelines establish when different species associate, but do not reveal which molecular regions form the interfaces. Therefore, we calculated the region-level intermolecular contact maps for each system (Figure 2). The functional A*β*_42_ regions were defined as metal-binding (residues 1–16), CHC (residues 17–21), polar (residues 22–29), and C-terminus, residues 30–42. For *α*-syn, the regions were defined as follows: N-terminus (residues 1–60), NAC (non-amyloid component, residues 61–95), and C-terminus (residues 96–140). The values in these heatmaps represent the cumulative contact probabilities between residue groups normalized over the full simulation time. Because the analyzed systems are dynamic and intrinsically disordered, the absolute percentages are expected to be low. Therefore, we interpret the contact maps primarily through relative differences between regions and systems rather than as direct binding times, which are covered in the previous section.

**Figure 2.**
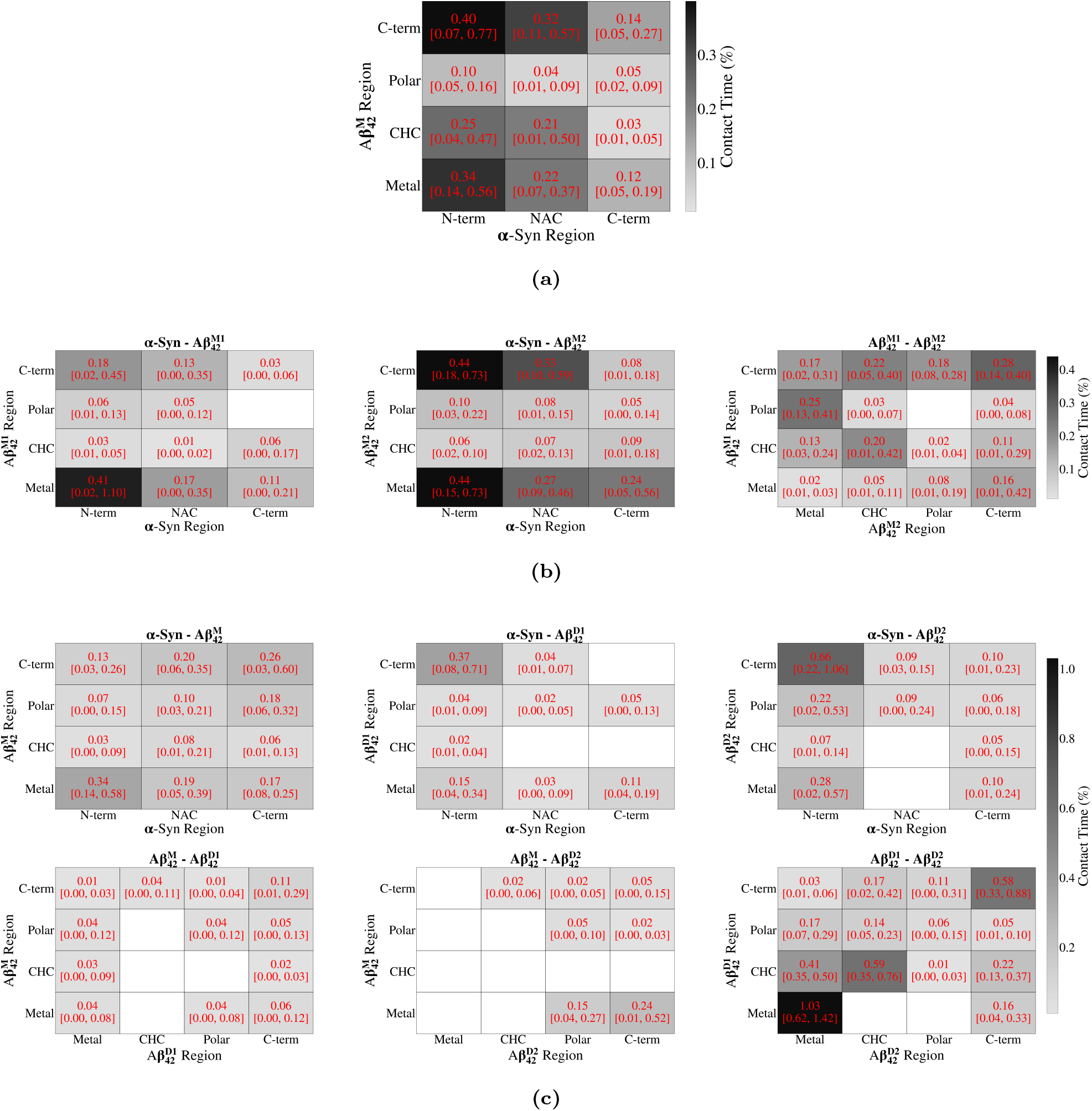
Polypeptide region contact heatmaps revealing binding interface preferences across simulation systems. **(a)** System 1: 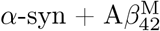. **(b)** System 2: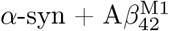 and 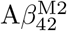. **(c)** System 3: 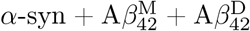 (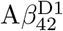 and 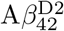 are labels distinguishing the two polypeptide chains within the dimer). A*β*_42_ regions: metal-binding (residues 1–16), CHC (residues 17–21), polar (residues 22–29), and C-terminus, residues 30–42. *α*-syn regions: N-terminus (residues 1–60), NAC (non-amyloid component, residues 61–95), and C-terminus, residues 96–140. The cell values represent the percentage of contact time between the specified polypeptide regions. Specifically, the values denote the cumulative sum of pairwise heavy-atom events within the specified regions, normalized against the total number of evaluated simulation frames and expressed as a percentage. The displayed values indicate the ensemble mean across all five simulation replicates. Bracketed values [*Lower, Upper*] report the 95% confidence interval, calculated via 1000 nonparametric bootstrap resamples of the independent replicates.

In System 1 (Figure 2a), the *α*-syn– 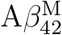 interface was dominated by contacts between the 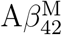 C-terminus, metal-binding region, and CHC with the *α*-syn N-terminal region. The 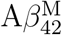 C-terminus also contacts the *α*-syn NAC domain. This contact pattern is consistent with a binding mode in which the N-terminal and NAC regions of *α*-syn engage both the charged and hydrophobic regions of A*β*_42_. Such an interface could partially shield the A*β*_42_ CHC/C-terminal surface from intramolecular coalescence or intermolecular A*β*_42_ contact. We examine this possibility further in Section 3.3, where we test whether *α*-syn engagement coincides with changes in A*β*_42_ *β*-hairpin formation.

In System 2 (Figure 2b), the presence of 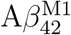 and 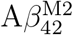 produced distinct inter-faces for heterogeneous and homogeneous contact. In *α*-syn– 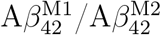 interactions,the *α*-syn N-terminus mainly engages the metal-binding region and C-terminus of A*β*_42_. Notably, the contact time between *α*-syn and the two A*β*_42_ monomers is heterogeneous: *α*-syn engages 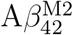 more strongly than 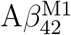 (for example, *α*-syn N-term × 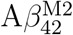 C-term = 0.44 vs. *α*-syn N-term × 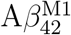 C-term = 0.18). This difference should not be interpreted as an intrinsic preference for one monomer over the other; rather, it reflects the different structural roles adopted by the two A*β*_42_ within the sampled trajectories. One monomer is more available for *α*-syn engagement, whereas the other contributes more to the A*β*_42_–A*β*_42_ interface.

The 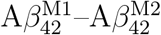 interface is stabilized by hydrophobic contacts involving the C-terminal and CHC regions of both peptides. Contacts between the 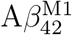 polar region and the 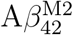 metal-binding region were also noted. The prominent participation of the C-terminus is consistent with the known role of the two additional hydrophobic residues in A*β*_42_, relative to A*β*_40_, in promoting early oligomerization [15]. The differ-ence observed in the 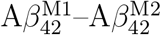 contact matrix is also consistent with the previous findings of Blinov et al. [65], showing that one chain contributes more strongly through the CHC and the other through the C-terminal region.

In System 3 (Figure 2c), the *α*-syn–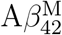 interface differed from that observed in System 1. The free 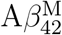 contacted *α*-syn mainly through its metal-binding region and C-terminus, whereas CHC showed fewer contacts. This indicates that, in the presence of a pre-formed A*β*_42_ dimer, the free monomer is engaged more peripherally and does not reproduce the CHC/C-terminal binding pattern observed for the isolated monomer in System 1.

The pre-formed dimer showed a stable internal 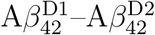 interface characterized by contacts among the metal-binding, CHC, and C-terminal regions of the two chains. This internal cohesion appears to reduce the accessibility of the dimer hydrophobic core to *α*-syn. Consequently, *α*-syn contacts with the CHC regions of the 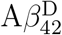 chains were limited in number. Instead, *α*-syn is bound mainly to exposed peripheral regions, including the C-termini and metal-binding regions of the dimer chains. This binding mode is consistent with the idea that *α*-syn can associate with an existing A*β*_42_ oligomeric surface without necessarily disrupting the internal dimer interface. This also agrees with reports suggesting that A*β*–*α*-syn interactions can occur through exposed oligomeric surfaces rather than through complete remodeling of pre-existing assemblies [27]. Therefore, System 3 should be interpreted as a comparison between one pre-organized A*β*_42_ interface and the initially unassociated monomeric systems, rather than as a general model of all pre-formed A*β*_42_ dimers.

This peripheral binding mode provides a possible atomistic basis for the experimentally observed A*β*–*α*-syn hybrid assemblies. AFM-based studies have identified distinct hetero-oligomer populations and suggested that A*β* and *α*-syn can co-localize through surface orientations that differ from those of homotypic assemblies [66, 67]. In the present simulations, the pre-formed A*β*_42_ dimer exhibits a distinct surface: its internal hydrophobic core remains engaged in A*β*_42_–A*β*_42_ contacts, while *α*-syn binds primarily to exposed peripheral regions. This suggests that the structural state of A*β*_42_ before *α*-syn encounter influences whether *α*-syn directly engages amyloidogenic CHC/C-terminal motifs or instead accommodates an already formed A*β*_42_ interface. This peripheral association provides an atomistic example of how *α*-syn may bind to an exposed A*β*_42_ oligomeric surface, as envisaged in models of heterotypic association and cross-seeding[3, 67]. Whether such an association subsequently promotes *α*-syn nucleation or cross-seeding cannot be determined from the present trajectories.

Across the three systems, A*β*_42_–*α*-syn contacts occurred mainly through the metal-binding region, CHC, and hydrophobic C-terminus of A*β*_42_ and the amphipathic N-terminal region and NAC domain of *α*-syn. These regions are consistent with the electrostatic and hydrophobic interactions expected to drive association: electrostatic contacts between the lysine-rich *α*-syn N-terminus and acidic residues in the A*β*_42_ N-terminal/metal-binding region, and hydrophobic contacts between the *α*-syn NAC domain and the CHC and C-terminal regions of A*β*_42_. The same interface components have been reported in previous computational studies of the A*β*_42_–*α*-syn heterodimer, where binding free-energy calculations attribute complex stability to both salt bridges involving the *α*-syn N-terminus and acidic A*β*_42_ residues and hydrophobic packing between the *α*-syn NAC and the A*β*_42_ C-terminal region [21, 22]; our contact maps recover these components together with contributions from the A*β*_42_ metal-binding region.

The dominance of the *α*-syn N-terminal and NAC regions in these interactions is consistent with prior studies showing that these domains contribute to both *α*-syn self-association and A*β*_42_ binding [34, 36]. Thus, the same molecular interfaces that participate in homomeric *α*-syn aggregation may also mediate heteromeric associations with A*β*_42_. This overlap suggests potential competition between *α*-syn self-association and A*β*_42_–*α*-syn co-assembly in A*β*-rich environments.

The intermolecular contact maps report each region-level pair as a trajectory average and therefore cannot reveal whether two pairs engage at the same moments in time. To resolve this, we computed the partial correlation between the per-frame contact counts of each of the two region-level pairs (Fig. S9), which shows that the pairs differ in their concerted engagement over time. Across all three systems, the correlation is predominantly intrasegmental: a single *α*-syn segment engages several A*β*_42_ regions whose contacts rise and fall together; thus, the segment acts as a unit. The clearest recurring motif is the concurrent engagement of the *α*-syn NAC with the A*β*_42_ CHC and polar regions — the cells (NAC – CHC) and (NAC – polar) — which is positive in every system (*r* = +0.28, +0.26, and +0.47 in Systems 1–3), with the CHC and polar regions being contiguous in the A*β*_42_ sequence. In System 2, the NAC instead engages the CHC and the metal-binding regions in an anticorrelated manner, the cells (NAC –CHC) and (NAC – metal-binding) being mutually exclusive (*r* = *−*0.25). Because the coefficients are resolved at the polypeptide rather than the individual species level, this exclusion may partly reflect the two A*β*_42_ monomers competing for the NAC rather than a single-interface conformational switch. A second recurrent motif couples the cells (C-terminus – C-terminus) and (C-terminus – metal-binding) in Systems 1 and 3 (*r* ≈ +0.26): this reports the same *α*-syn C-terminal segment engaging both ends of A*β*_42_ at once.

The systems differ mainly in the balance between concurrent and mutually exclusive engagement. System 1, in which *α*-syn faces a single A*β*_42_ monomer, is almost entirely concurrent (eight of ten reproducible couplings), consistent with the construction and maintenance of a single interface. System 2, where *α*-syn is presented with two independent A*β*_42_ monomers, carries the most mutually exclusive couplings (eight of seventeen), consistent with a single *α*-syn chain having to partition its contacts between competing partners rather than engaging one interface. Its strongest observed correlation is nonetheless concurrent and uniquely couples two *different α*-syn segments: engagement of the N-terminus correlates with that of the NAC, such that when one segment binds, the other tends to be bound at the same time. System 3, with a pre-formed A*β*_42_ dimer, returns to predominantly concurrent engagement (11 of 15) and shows the strongest single coordination of the study within the NAC, coupled with adjacent CHC and polar regions (*r* = +0.47). Throughout, the joint-occupancy *p*_joint_ shows that strongly correlated coupling does not correspond to a dominant state: the recurring NAC–CHC/NAC–polar motif, though the most reproducible, is occupied in only about a third of engaged frames (*p*_joint_ = 0.34), marking it a well-defined but transiently sampled binding mode, whereas the System 2 N-terminus/NAC coupling with A*β*_42_ C-terminus/metal-binding is both highly correlated and a dominant configuration (*p*_joint_ = 0.88).

Together, Sections 3.1 and 3.2 show that *α*-syn associates with all A*β*_42_ species studied here, but with distinct persistence patterns, interface preferences, and internal coordination. Across all assembly states, its contacts are organized rather than independent: individual *α*-syn segments engage several A*β*_42_ regions as temporally correlated units, which converge preferentially on the CHC and C-terminus. With monomeric or nascent dimeric A*β*_42_, *α*-syn frequently contacts these same nucleation-prone regions, whereas in the presence of a pre-formed A*β*_42_ dimer, it binds more peripherally, while the internal dimer interface remains intact. Because the positively correlated contacts converge on the CHC and C-terminal regions that drive A*β*_42_ nucleation, they raise the next mechanistic question: do these binding modes alter A*β*_42_ conformational maturation, particularly the formation of CHC/C-terminal *β*-hairpin motifs associated with amyloid-prone monomeric conformations [15, 63]?

### 6.3 Structural consequences of α-syn–Aβ_42_ aggregation

The formation of a stable *β*-hairpin within the A*β*_42_ monomer is widely considered to be the rate-limiting step in amyloid nucleation, oligomer growth, and fibril elongation [13]. To determine whether *α*-syn promotes or suppresses these amyloidogenic motifs, we quantified region-specific *β*-hairpin occurrence across all replicates for 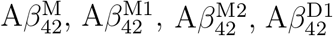,and 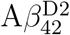 (Figure 3). In this analysis, a *β*-hairpin was counted only when the motif involved the CHC and/or C-terminal region, thereby focusing on folding events most directly related to A*β*_42_ nucleation. Furthermore, a comparison of the three systems allowed us to test whether the binding modes identified above correlated with the promotion, suppression, or preservation of the intramolecular folding motifs associated with early amyloidogenic maturation.

**Figure 3.**
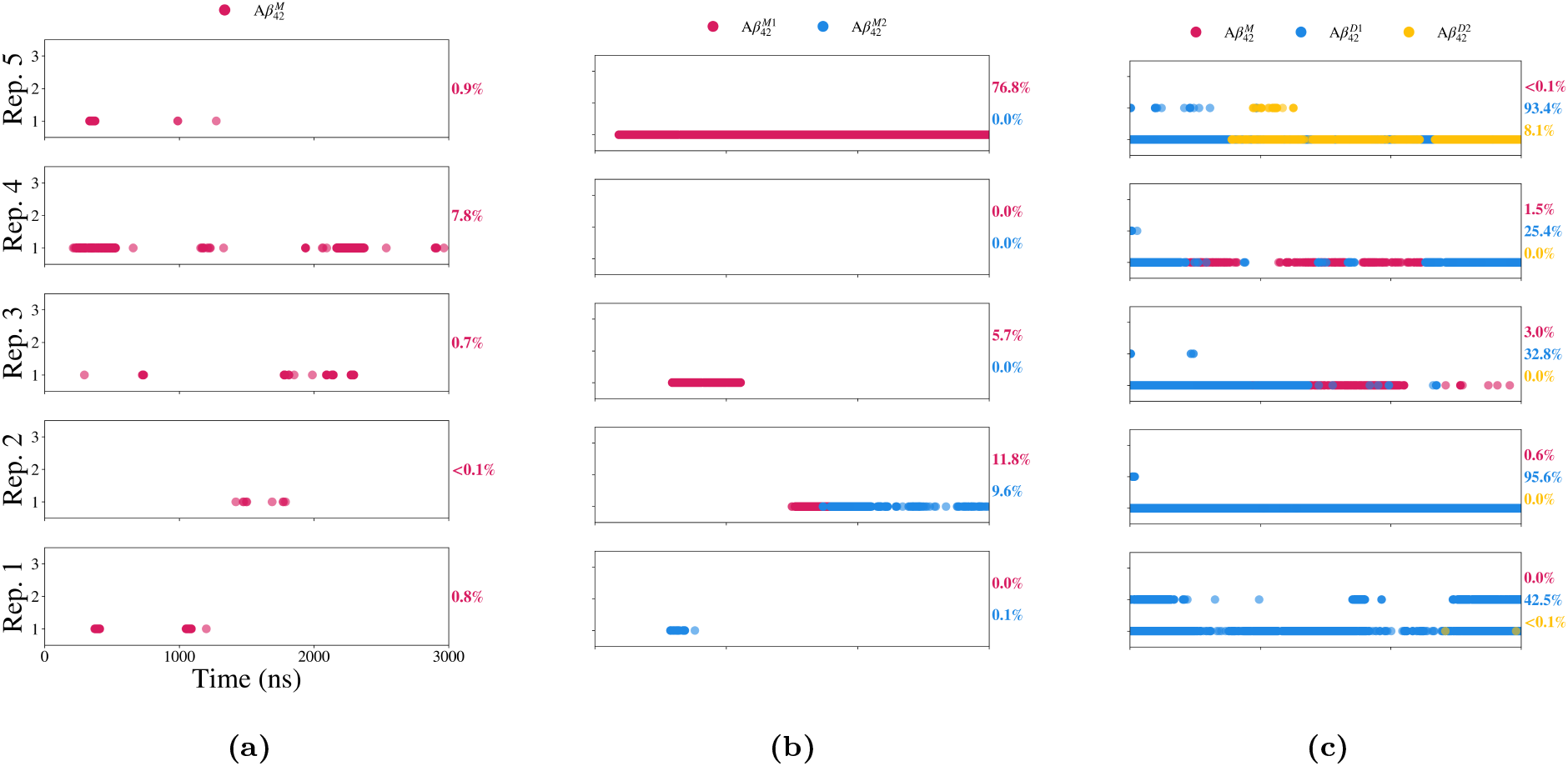
*β*-Hairpin formation analysis. Temporal evolution of *β*-hairpin counts for **(a)** System 1: 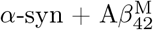. **(b)** System 2: 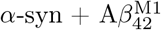 and 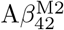. **(c)** System 3: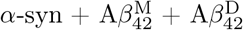 (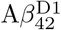 and 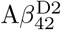). The percentages on the right vertical axis indicate the total proportion of time each respective A*β*_42_ chain successfully maintained a region-specific *β*-hairpin (CHC/C-terminus) during that individual 3000 ns replicate.

In System 1 (Figure 3a), *β*-hairpin formation remained sparse and transient despite the frequent formation of the *α*-syn–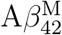 heterodimer. In most replicates, hairpin motifs rarely persisted beyond a few nanoseconds and occupied less than 1% of the trajectory. Replicate 2 provides a representative example: the heterodimer remained aggregated for 95% of the simulation, whereas the *β*-hairpin occupancy was below 0.1%. Contact analysis showed that the *α*-syn NAC region engaged both the 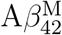 CHC and C-terminus, with cumulative contact values of 0.81% and 0.79%, respec-tively (SI Figure S5 (a)). Replicates 1, 3, and 5 exhibited similar inverse relationships. In these trajectories, the polypeptides remained associated for 65%, 75%, and 80% of the simulation, respectively, while the *β*-hairpin occupancy remained below 1%. The corresponding contact maps indicate that the *α*-syn N-terminal region frequently engaged the 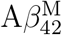 CHC and C-terminus (SI Figures S4a, S6a, and S8a).

These observations suggest that the occupation of the A*β*_42_ CHC/C-terminal surface by the *α*-syn N-terminal or NAC regions reduces the conformational freedom required for intramolecular *β*-hairpin formation. The observations in Replicate 4 lend further credence to this conclusion. In this trajectory, *α*-syn– 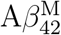 aggregation oc-curred for only 16% of the simulation, whereas the *β*-hairpin occupancy reached 7.8% of the simulation. Consistent with this behavior, the 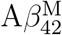 C-terminus showed minimal contact with *α*-syn, with a cumulative contact value of 0.04%, and the CHC was not detected (SI Figure S7a). Thus, the reduced occupation of the CHC/C-terminal surface coincided with increased sampling of hairpin-prone conformations.

The hairpin topologies of System 1 (SI Table S1) indicate that the residual folding events did not converge on a single dominant structural register. Together, these results are consistent with a competitive mode of molecular crosstalk in which *α*-syn association does not prevent heterodimer formation but limits the intramolecular ordering of A*β*_42_. This interpretation agrees with experimental observations that *α*-syn can inhibit A*β*_42_ fibrillization while redirecting A*β*_42_ toward soluble oligomeric assemblies [26]. However, the present simulations characterized early structural events and did not directly determine the long-term fibrillization outcome or cytotoxicity of the resulting complexes.

System 2 revealed a more heterogeneous relationship between A*β*_42_ homodimers, *α*-syn binding, and *β*-hairpin formation (Figure 3b). In replicate 5, *α*-syn showed negligible contact with either A*β*_42_ chain, with all regional contact values below 0.01%. This allowed the 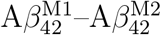 association to proceed with minimal heterotypic interference. Formation of the stable A*β*_42_ homodimer coincided with a persistent *β*-hairpin signal localized primarily to 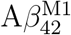.

The contact map showed an asymmetric homodimer interface (SI Figure S8b). The strongest contacts occurred between the 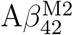 C-terminus and the 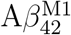 metal-binding region, supplemented by C-terminus–C-terminus contacts, whereas the CHC regions of both chains remained comparatively weakly engaged. Although both chains contributed C-terminal residues to the interface, 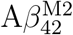 made a larger intermolecular C-terminal contribution. Residue-level analysis showed that the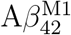 hairpin adopted a C-term ↔C-term topology (SI Table S1). This topology resembled the dominant motif observed for 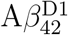 in four of the five System 3 replicates, although the strand registers were different. The 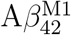 hairpin was more compactly localized within residues 32–37, whereas the 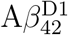 hairpins generally extended through residues 35–42. Glycine residues frequently occur within the intervening turns, consistent with their conformational flexibility.

In contrast, replicate 4 captured 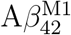 and 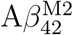 forming a nascent homodimer that was extensively engaged by *α*-syn (Figure 1b), and neither A*β*_42_ chain formed a detectable region-specific *β*-hairpin. The corresponding contact map showed that the *α*-syn N-terminal and NAC regions contacted the 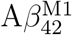 metal-binding region and the C-termini of both A*β*_42_ chains (SI Figure S7b). Simultaneously, the A*β*_42_ chains formed substantial intermolecular CHC–CHC, CHC–C-terminal, and C-terminal–C-terminal contacts, with cumulative values of 0.42%, 0.46%, and 0.36%, respectively.

This contact pattern suggests that simultaneous *α*-syn engagement and strong A*β*_42_ intermolecular interactions constrain the CHC and C-terminal regions, reducing the conformational freedom required for intramolecular hairpin formation. Rather than demonstrating that *α*-syn physically prevents folding, the trajectory supports a model in which *α*-syn stabilizes an alternative nascent dimer interface that is less compatible with hairpin motifs. This interpretation is consistent with the proposal that *α*-syn can redirect A*β*_42_ toward non-fibrillar oligomeric states [26].

Replicates 1 and 3 exhibited similar behaviors (Figure 1b). In both trajectories, an 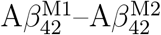 dimer formed and subsequently remained associated with *α*-syn for most of the simulation. Although the magnitudes of the regional contacts differed from replicate 4, both trajectories showed low hairpin occupancies for the A*β*_42_ chains (Figure S4b and Figure S6b). In replicate 3, 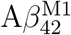 was less extensively engaged by *α*-syn than in replicate 1 and retained a measurable hairpin occupancy of 5.7%, whereas no corresponding motif was detected in replicate 1. In both trajectories, the more strongly engaged 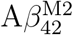 chain showed little or no hairpin formation.

Replicate 2 provides an important qualification to the association observed in the remaining trajectories. Although *α*-syn remained associated predominantly with 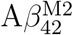 for approximately 80% of the simulation (Figure 1b), both A*β*_42_ chains sampled region-specific hairpins with occupancies of 11.8% and 9.6%. Therefore, contact persistence alone is not predictive of hairpin suppression. A long-lived interface may leave the intramolecular CHC/C-terminal degrees of freedom required for hairpin formation partially accessible, particularly when the contact is distributed over peripheral or non-competing regions. In contrast, the low hairpin occupancies in replicates 1, 3, and 4 coincided more closely with interfaces involving the CHC and C-terminal regions and, in some cases, with simultaneous A*β*_42_–A*β*_42_ contacts. Therefore, the comparison supports a geometry-dependent association between the interface composition and hairpin sampling, rather than a general inhibitory effect of *α*-syn binding. Collectively, the System 2 trajectories indicate that *α*-syn association can bias the organization of a nascent A*β*_42_ dimer interface. Whether this bias coincides with reduced *β*-hairpin sampling depends on which regions are engaged and how heterotypic and homotypic contacts are spatially combined. The trajectories do not provide a deterministic criterion for separating the inhibitory and noninhibitory contact geometries.

In System 3 (Figure 3c), 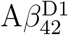 displayed persistent region-specific *β*-hairpin signatures across all five replicates. In contrast, 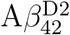 and the free 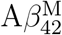 showed primarily short and transient hairpin events, generally occupying less than 5% of the trajectory,with the exception of an 8.1% occupancy for 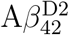 in replicate 5. Contact analysis revealed that the 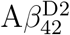 C-terminus contributed more extensively to the intermolecular dimer interface than the corresponding region of 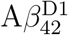 (Figure 2c). This difference parallels the behavior observed in System 2 replicate 5: the more strongly committed 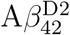 C-terminus has less conformational freedom for intramolecular folding, whereas the less constrained 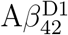 chain can form persistent hairpin motifs.

The dominant 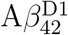 folding motif was a C-term*↔*C-term hairpin in four of the five replicates. A representative register contained *β*-strand segments at residues 36–37 and 40–41, separated by a turn involving residues 38–39 (SI Table S1). Replicate 4 was the exception, with 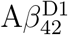 adopting a CHC*↔*C-terminal register spanning approximately residues 20–32. These results indicate that the pre-formed dimer supports persistent intramolecular ordering within a chain while retaining a structurally asymmetric intermolecular interface.

As shown by the contact maps, *α*-syn associated predominantly with exposed peripheral regions of the pre-formed dimer and made comparatively few contacts with its CHC regions. The persistence of the 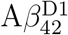 hairpin across contact states suggests that peripheral *α*-syn binding does not substantially disrupt the internal structure of this particular pre-formed dimer. Within the accessible trajectories, *α*-syn associated predominantly with exposed peripheral regions, while no substantial remodeling of the buried interface of this particular A*β*_42_ dimer was detected. This observation is compatible with a non-disruptive peripheral association [36]. However, it does not distinguish passive non-interference from the active stabilization of the dimer.

Hairpin formation in the free 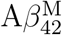 of System 3 remained intermittent throughout the simulation. This behavior is compatible with a pathway in which intermolecular association precedes slower conformational ordering, as described by the “dock-and-lock” models of amyloid growth [68]. However, because no complete elongation event was sampled here, the observed fluctuations should be interpreted as early conformational rearrangements rather than direct evidence of a full dock-and-lock process.

To determine whether these changes in region-specific *β*-hairpin formation were accompanied by broader conformational changes, we compared the contact-dependent structural ensembles of the A*β*_42_ chains (Figure 4A). Across the monomeric A*β*_42_ chains, contact with *α*-syn generally shifted the radius-of-gyration distribution toward larger values and increased their breadth. These changes indicate a greater overall conformational extension and heterogeneity in the *α*-syn-associated ensembles.

**Figure 4.**
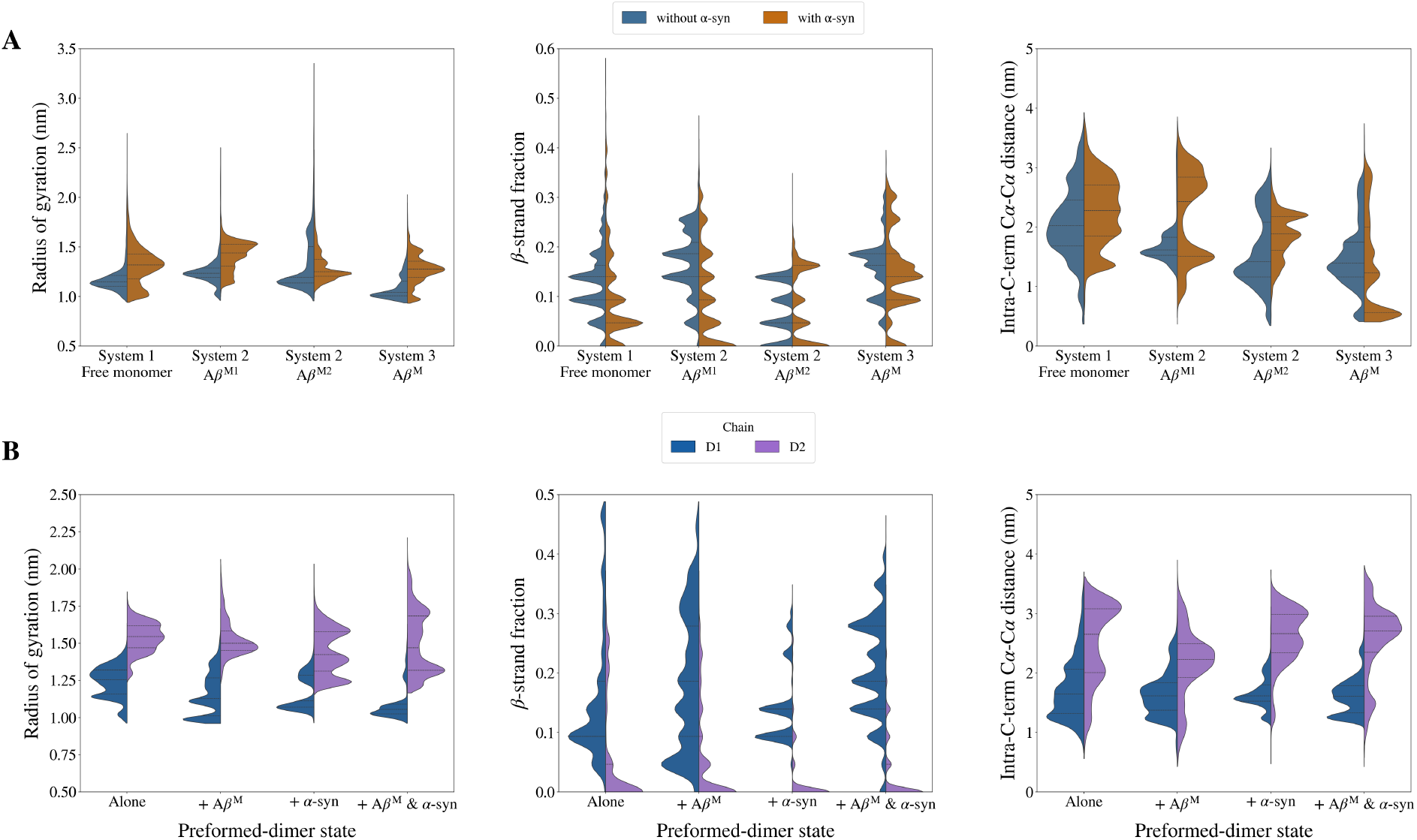
Structural ensemble distributions of A*β*_42_ chains as a function of *α*-syn contact state, across all three simulated systems. Split violins show the per-frame distribution of three structural metrics — radius of gyration (*R*_g_), *β*-strand fraction, and intra-C-terminal C*α*–C*α* distance (residues 30–42) — partitioned by the absence (blue) or presence (orange) of *α*-syn contact, as defined by the dual-criteria aggregation metric. **(A)** Free A*β*_42_ monomer chains across Systems 1, 2, and 3. **(B)** Pre-formed A*β*_42_ dimer chains D1 and D2 (System 3) across four contact states: alone, in contact with 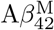, in contact with *α*-syn, and in contact with both 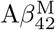 and *α*-syn. Each violin pools frames across five independent 3 µs replicates from each system.

The exception was 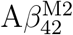 in System 2, for which the median *R*_g_ remained similar between the contact and non-contact states. This limited shift may reflect the heterogeneous binding geometries sampled by 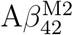 simulations. Although *α*-syn frequently contacts its metal-binding and C-terminal regions, these interactions do not produce a consistent directional change in global chain dimensions. Thus, engagement by *α*-syn can alter the local A*β*_42_ structure without necessarily causing uniform compaction or extension of the entire chain.

The *β*-strand fraction was generally lower when the monomeric A*β*_42_ chains were in contact with *α*-syn. This trend parallels the reduction in region-specific *β*-hairpin formation described above. Although the *β*-strand fraction reports the global secondary-structure content of each chain rather than the CHC/C-terminal motif specifically, the concurrent decrease in both quantities suggests that *α*-syn engagement shifts the A*β*_42_ monomer ensemble away from ordered conformations associated with early amyloidogenic maturation.

Consistent with this interpretation, the intra-C-terminal C*α*–C*α* distance between residues 30 and 42 generally increased upon *α*-syn contact. A larger separation indicates that compact C-terminal conformations associated with hairpin nucleation [15] were sampled less frequently in the *α*-syn-associated ensemble. An exception was observed for the free 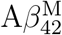 in System 3, for which the intra-C-terminal distance did not increase appreciably upon *α*-syn-contact. This limited response is consistent with the binding pattern identified in Section 3.2, where *α*-syn interacted preferentially with exposed regions of the pre-formed 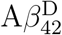 chains and engaged the free monomer less extensively than in System 1.

System 2 shows two distinct sub-populations in the intra-C-terminal distance upon *α*-syn contact. The lower-distance peak (~ 1.5 nm) corresponds to frames from replicate 5, in which 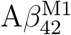 and 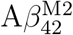 formed a nascent homodimer that was not engaged by *α*-syn, and 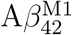 showed a sustained *β*-hairpin signal. The higher-distance peak reflects the remaining replicates, where *α*-syn contact with individual monomers results in a more extended C-terminal conformation.

The pre-formed dimer chains exhibited distinct structural behaviors across all three metrics (Figure 4B). 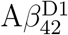 consistently presents a lower *R*_g_, a higher *β*-strand fraction, and a shorter intra-C-terminal C*α*–C*α* distance compared to 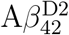 across all four contact states. The latter two metrics are in direct agreement with the persistent *β*-hairpin signal observed for 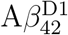 across all five System 3 replicates (Figure 3c), where the C-terminal of 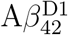 preserves sufficient conformational freedom for intramolecular nucleation owing to lower intermolecular commitment relative to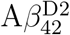.

Importantly, the *β*-strand fraction and intra-C-terminal distance of 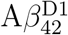 remained comparatively stable across different contact states. This suggests that contact with either *α*-syn or the free 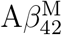 does not substantially perturb the structural features retained by this chain within the pre-formed dimer. This observation is consistent with the peripheral binding mode described in Section 3.2, in which *α*-syn primarily engages the exposed regions of the dimer without extensively contacting its buried CHC-rich interface. Within the sampled trajectories, this pattern is most conservatively described as peripheral association without detectable disruption of the characteristic structural features of the selected dimers. [36].

In contrast to the contact-dependent changes observed for the A*β*_42_ monomer ensembles, the global structural properties of *α*-syn were less affected by A*β*_42_ association (Figure S10). In Systems 1 and 2, the distributions of *R*_g_, *α*-helix fraction, and NAC-to-C-terminal C*α*–C*α* distance showed no large or consistent shifts between the contact states. The *α*-helix fraction was also similar across the analyzed states, providing no evidence of a substantial global helix-inducing effect of A*β*_42_ binding under these simulation conditions.

An exception was observed for System 3, where the NAC-to-C-terminal distance distribution shifted toward lower values upon A*β*_42_ contact. This shift suggests partial compaction of the NAC–C-terminal geometry when *α*-syn associates with the preformed dimer. However, the magnitude of this change was smaller than that of the contact-dependent shifts observed for the A*β*_42_ monomer chains. Thus, among the structural metrics analyzed here, *α*-syn association perturbs the global conformational ensemble of monomeric A*β*_42_ more strongly than A*β*_42_ association perturbs *α*-syn. The heteroptic association can be seen in the video clip “Heterotipic association clip” (SI). The clip starts with the early association of all species in System 3 and shows its evolution for approximately 2500 frames. The polypeptide regions are colored from lighter to darker shades: lighter, the N-Terminus/metal-binding and darker, the C-terminus regions.

Taken together, the ensemble-level distributions support an oligomeric-state-dependent response to *α*-syn binding. Contact with *α*-syn generally shifts monomeric A*β*_42_ toward more extended conformations with reduced *β*-strand and *β*-hairpin content, whereas the characteristic structural features of the pre-formed A*β*_42_ dimer remain comparatively stable. These results are consistent with a mechanism in which *α*-syn perturbs lower-order A*β*_42_ species more readily than it remodels an already established dimeric interface.

To evaluate whether these structural shifts were accompanied by broader changes in the backbone conformational sampling, we calculated the first-order Shannon entropy, *S*_1_, from the residue-wise backbone dihedral angle distributions (*ϕ* and *ψ*) (Figure S12). This analysis compared the local dihedral sampling in the free and aggregated states. Because *S*_1_ is calculated from individual dihedral distributions and does not include correlations between residues, it should be interpreted as a measure of local conformational diversity rather than a complete thermodynamic conformational entropy.

Across the A*β*_42_ species, the first-order dihedral entropy *S*_1_ differed only modestly between the free and associated states. This observation does not imply that the structural ensembles are not altered. The radius of gyration, *β*-strand fraction, and intra-C-terminal distance report shifts in selected collective features, whereas *S*_1_ measures the summed marginal diversity of individual backbone dihedrals and does not include correlations between them. Therefore, a redistribution among extended, compact, or *β*-structured conformations can alter these collective observables without producing a comparable reduction in the total marginal dihedral diversity.

The combined results suggest that *α*-syn association biases the relative populations of particular A*β*_42_ conformations, especially those involving C-terminal compaction and *β*-structure, rather than globally immobilizing the backbone. A*β*_42_ consequently remains locally heterogeneous even when the associated ensemble is, on average, more extended and samples fewer *β*-hairpin-like states. Because *S*_1_ excludes inter-coordinate correlations and does not separate changes in accessible states from changes in their exploration rates, it was used here only as a descriptive measure of local conformational diversity.

The corresponding *S*_1_ values for *α*-syn also remained close to those in the free state. Within the resolution of this first-order metric, heterotypic association did not produce a global loss of local backbone diversity, although more specific correlated or domain-level rearrangements cannot be excluded.

## 4. Conclusion

The reported simulations characterized early, configuration-dependent encounter complexes between *α*-syn and three selected A*β*_42_ arrangements. In systems initiated from monomeric A*β*_42_, *α*-syn frequently contacted the metal-binding, CHC, and C-terminal regions. In several trajectories, engagement of the CHC and C-terminal regions coincided with reduced *β*-hairpin occupancy, larger radii of gyration, increased intra-C-terminal distances, and lower *β*-strand fractions. These relationships were not uniform across the replicates, demonstrating that contact duration alone is insufficient to predict the structural response; the geometry and regional composition of the interface are also important. Moreover, these contacts were temporally coordinated, with individual *α*-syn segments engaging several A*β*_42_ regions as concerted units — most reproducibly the NAC with the contiguous CHC and polar regions. The balance between concurrent and mutually exclusive engagement differed with the A*β*_42_ configuration, shifting toward mutually exclusive engagement when two independent monomers competed for a single *α*-syn chain.

In the system containing a pre-formed A*β*_42_ dimer, *α*-syn interacted mainly with exposed peripheral regions, while the characteristic *β*-structured features of this particular dimer remained comparatively stable. The simulations therefore reveal a contrast between the greater contact-dependent responsiveness of monomeric A*β*_42_ and the relative structural persistence of the selected pre-formed dimer. They did not establish that peripheral *α*-syn binding actively stabilizes the dimer or that the same behavior occurs for other dimer polymorphs.

First-order dihedral entropy analysis further indicated that these changes in the selected collective structural features were not accompanied by a global loss of the local backbone diversity. The association appears to redistribute the populations of particular conformational states while preserving substantial local heterogeneity.

Overall, the results support a model in which the early structural consequences of *α*-syn–A*β*_42_ association depend on the initial A*β*_42_ configuration and the detailed interface formed in each encounter. Terms such as shielding- and scaffold-like associations provide useful descriptions of the limiting behaviors observed; however, the present data do not define a universal mechanistic boundary between them.

The scope of these conclusions is limited by the exploratory design of the study. The simulations covered three selected molecular compositions, five trajectories per system, and one pre-formed A*β*_42_ dimer geometry over 3 *µ*s timescales. They do not sample the full distribution of A*β*_42_ oligomeric states, higher-order oligomers, alternative stoichiometries, concentration-dependent effects, or physiological A*β*_42_/*α*-syn ratios. Moreover, the first-order entropy approximation excludes correlations between the internal coordinates. Therefore, the reported patterns should be interpreted as comparative observations within the simulated systems and as hypotheses for subsequent testing rather than as universal mechanisms of A*β*_42_–*α*-syn co-aggregation.

## Declaration of generative AI and AI-assisted technologies in the writing process

The authors used ChatGPT exclusively for language editing to improve the clarity and readability of the manuscript. Gemini was used to assist in generating Python and Bash scripts for data analysis. All codes were verified and validated by the authors. No generative AI tools were used for data analysis, interpretation, or drafting of the scientific content. The authors are fully responsible for all content and conclusions presented.

## Data Availability

The solvent-stripped simulation trajectories for the aqueous systems (water and ions removed; PBC-corrected with `autoimage` and `fiximagedbonds`, centered on the first A*β*_42_ chain, and frame-by-frame RMS aligned on its C_*α*_ atoms, rms:1-43@CA previous) are available at DOI:10.5281/zenodo.21930581. The repository also contains the analysis scripts (trajectory analysis and convergence diagnostics) and the input and topology files needed to reproduce the simulations.

## Acknowledgments

This work was completed with resources provided by the University of Coimbra CERES Research Center, which is supported by FCT, within the projects DOI: 10. 54499/UID/00102/2025 (Base and Programmatic funding) and DOI: 10.54499/UID/ PRR/00102/2025.

