## Supplementary Information for "Beyond Co-Existence: Early *α*-Synuclein-Amyloid-*β*_42_ Association from Molecular Dynamics Simulations"

### Supporting Information

#### Beyond Co-Existence: $\alpha$-Synuclein Sequesters Amyloid-$\beta_{42}$ into Distinct Hybrid Assemblies — A Molecular Dynamics Study

**F. Carvalho**[1], **P. Maximiano**[2], **M. Hashemi**[2,*], **P. N. Simões**[1,*]

[1]University of Coimbra, CERES, Department of Chemical Engineering,

Rua Sílvio de Lima, Coimbra, 3030-790, Portugal

[2]Department of Physics, Auburn University,

Leach Science Center 3126, Auburn, AL, 36849-5319, USA

### I Convergence Diagnostics

The convergence diagnostics below characterize the breadth of sampling accessible within each 3 µs replicate. For each system and replicate, we report three standard diagnostics for the $\alpha$-syn/A$\beta_{42}$-monomer interaction and for the radius of gyration of each chain: **(A)** block averages over ten equal blocks, **(B)** the cumulative running average, and **(C)** the temporal autocorrelation function.

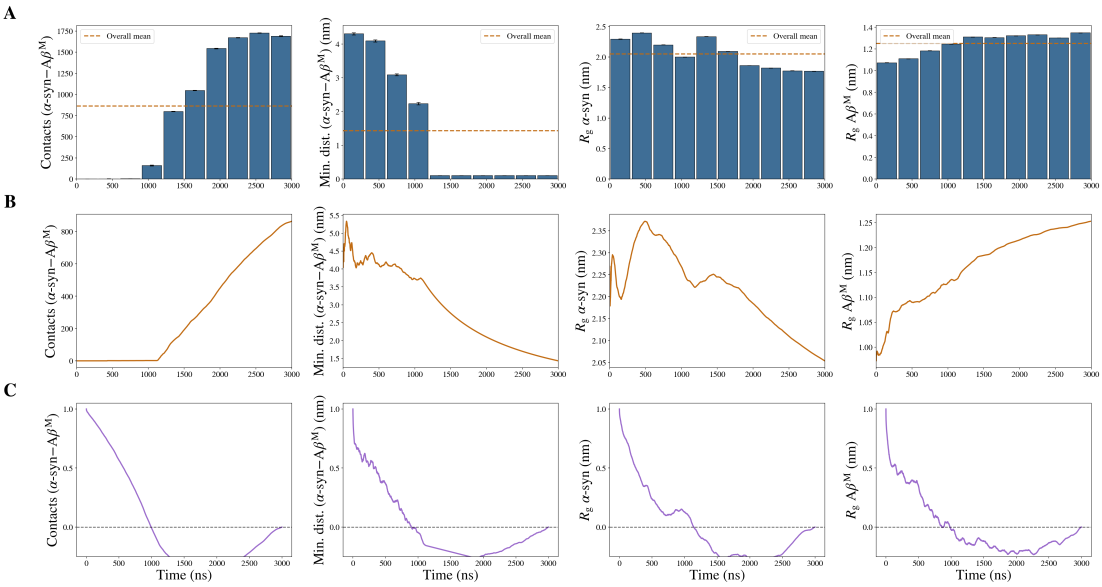
A
B
C
Overall mean
Contacts (α-syn–Aβ^M)
Min. dist. (α-syn–Aβ^M) (nm)
R_g α-syn (nm)
R_g Aβ^M (nm)
Time (ns)

(a)

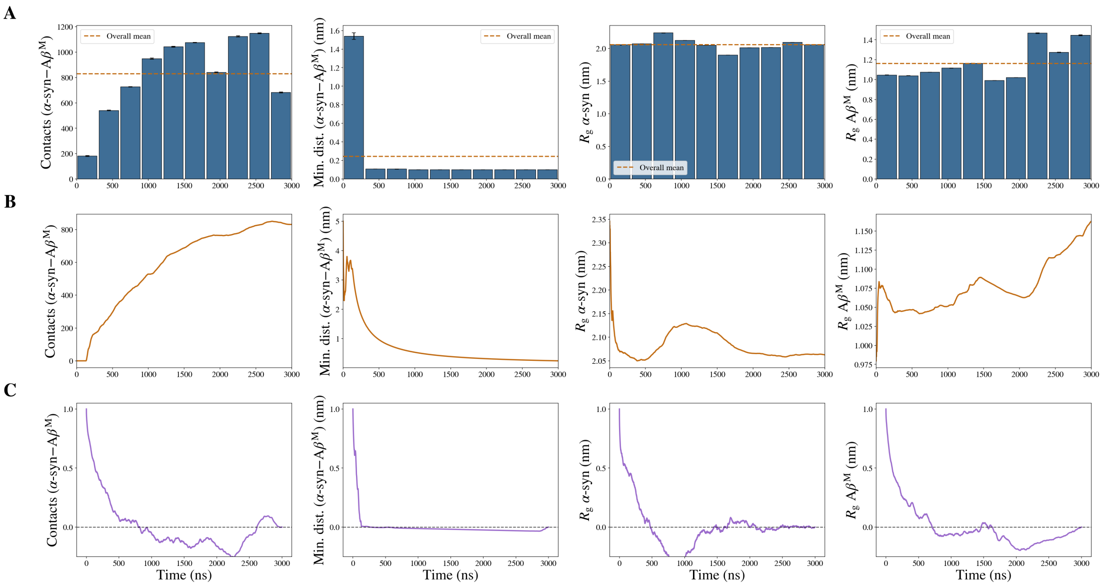
A
B
C
Overall mean
Contacts (α-syn–Aβ^M)
Min. dist. (α-syn–Aβ^M) (nm)
R_g α-syn (nm)
R_g Aβ^M (nm)
Time (ns)

(b)

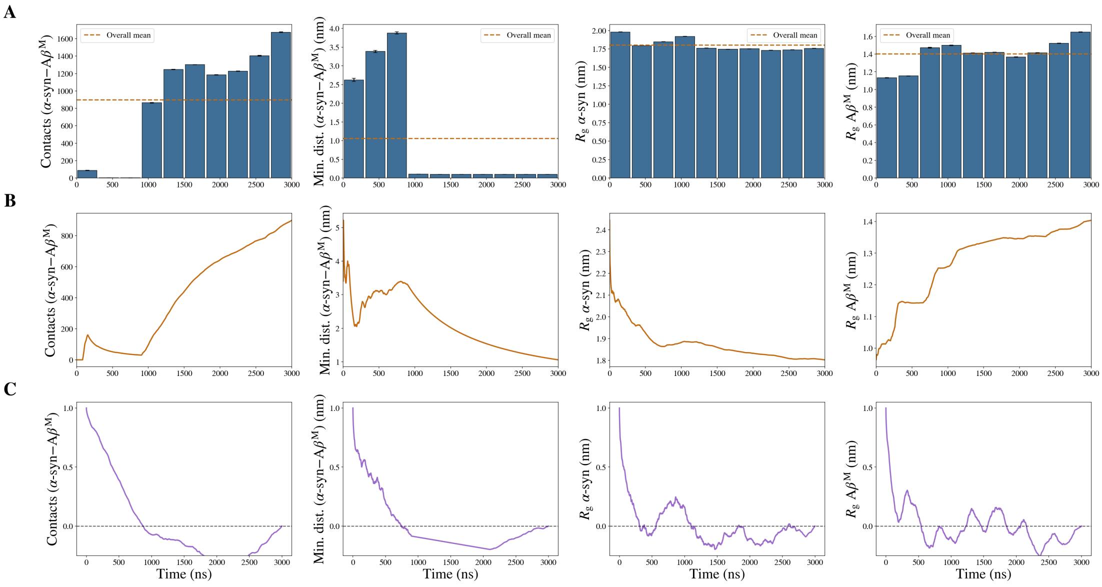
A
B
C
Overall mean
Contacts (α-syn–Aβ^M)
Min. dist. (α-syn–Aβ^M) (nm)
R_g α-syn (nm)
R_g Aβ^M (nm)
Time (ns)

(c)

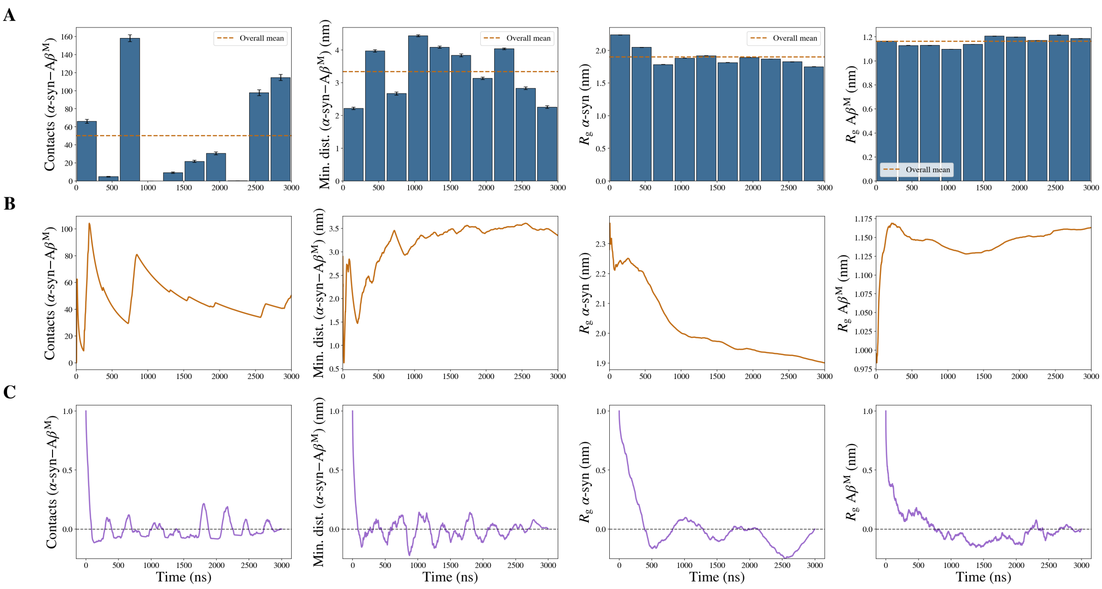


**(d)**

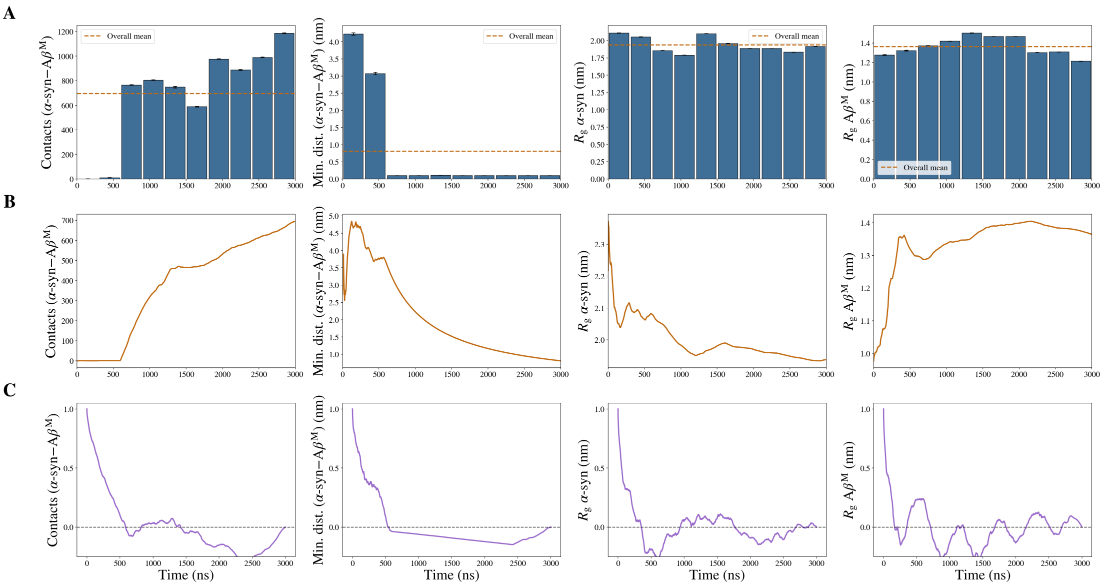


**(e)**

**Figure S1.** Sampling diagnostics for **System 1** ($\alpha$-syn + A$\beta_{42}^{\mathrm{M}}$), replicates 1–5 (panels a–e). Within each panel, columns show (left to right) the inter-chain heavy-atom contacts, minimum inter-chain distance, $\alpha$-syn radius of gyration, and A$\beta_{42}^{\mathrm{M}}$ radius of gyration. Rows show **(A)** block averages over ten equal blocks (error bars: standard error of the mean within each block; dashed line: overall trajectory mean), **(B)** the cumulative running average, and **(C)** the temporal autocorrelation function.

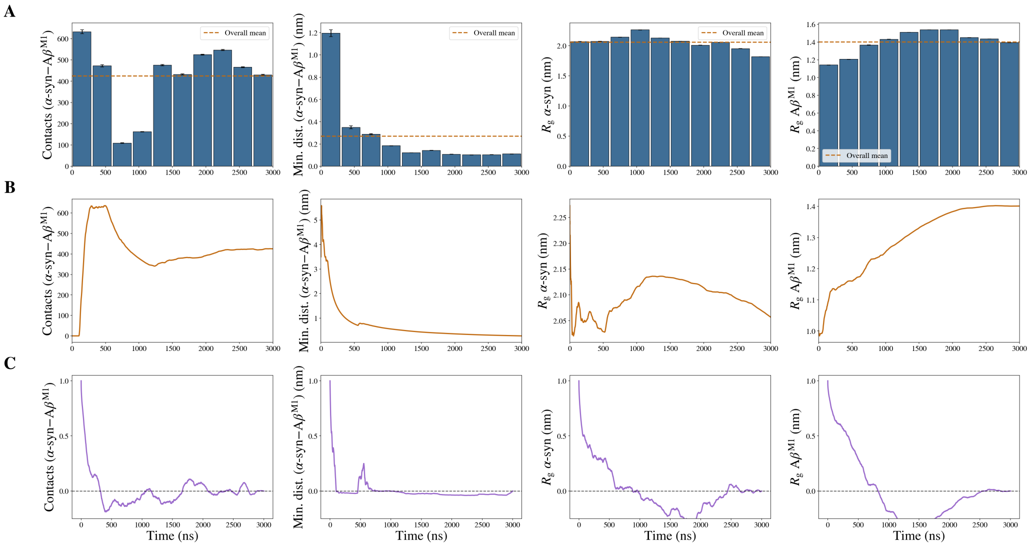
A
B
C
Overall mean
Contacts (α-syn–Aβ$^{M1}$)
Min. dist. (α-syn–Aβ$^{M1}$) (nm)
$R_g$ α-syn (nm)
$R_g$ Aβ$^{M1}$ (nm)
Time (ns)

(a)

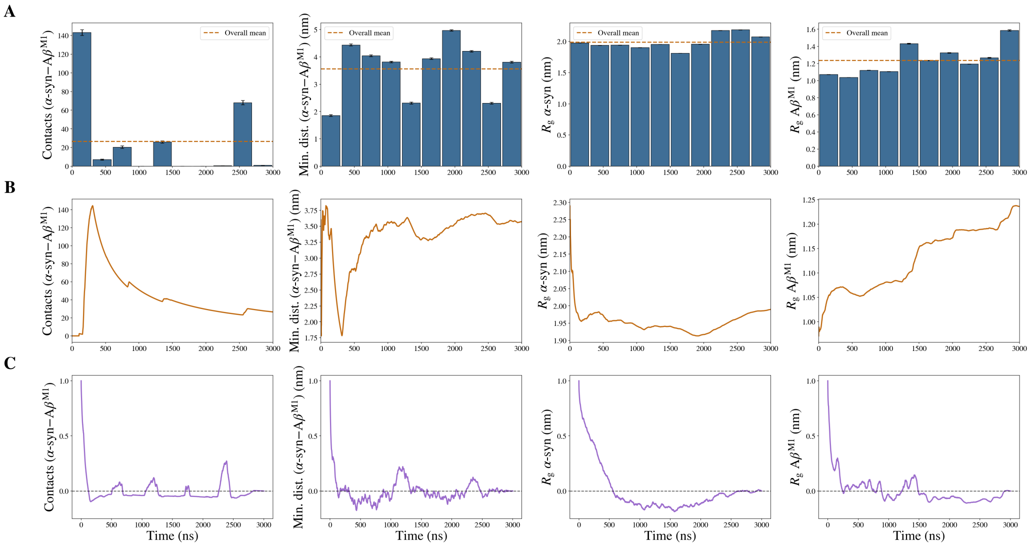
A
B
C
Overall mean
Contacts (α-syn–Aβ$^{M1}$)
Min. dist. (α-syn–Aβ$^{M1}$) (nm)
$R_g$ α-syn (nm)
$R_g$ Aβ$^{M1}$ (nm)
Time (ns)

(b)

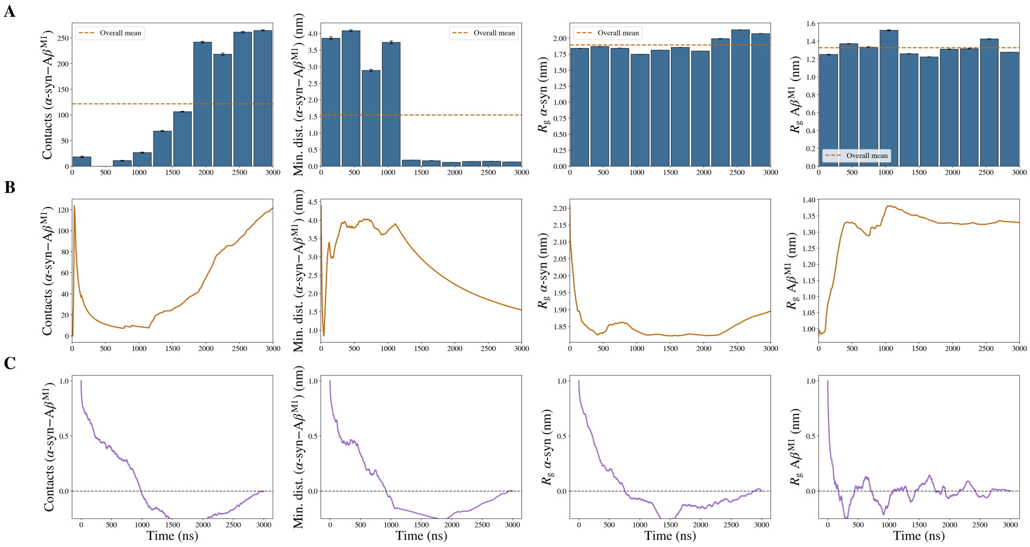
A
B
C
Overall mean
Contacts (α-syn–Aβ$^{M1}$)
Min. dist. (α-syn–Aβ$^{M1}$) (nm)
$R_g$ α-syn (nm)
$R_g$ Aβ$^{M1}$ (nm)
Time (ns)

(c)

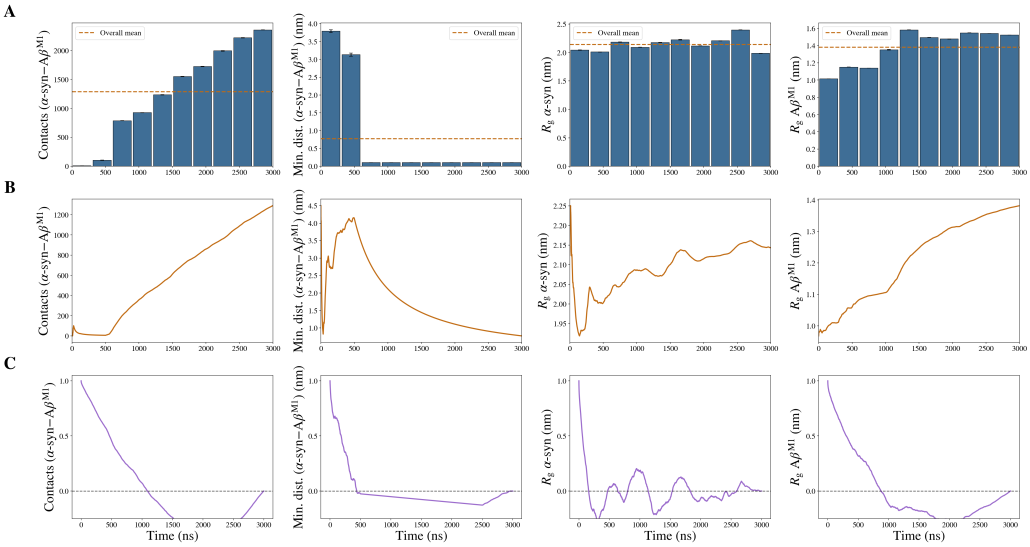


**(d)**

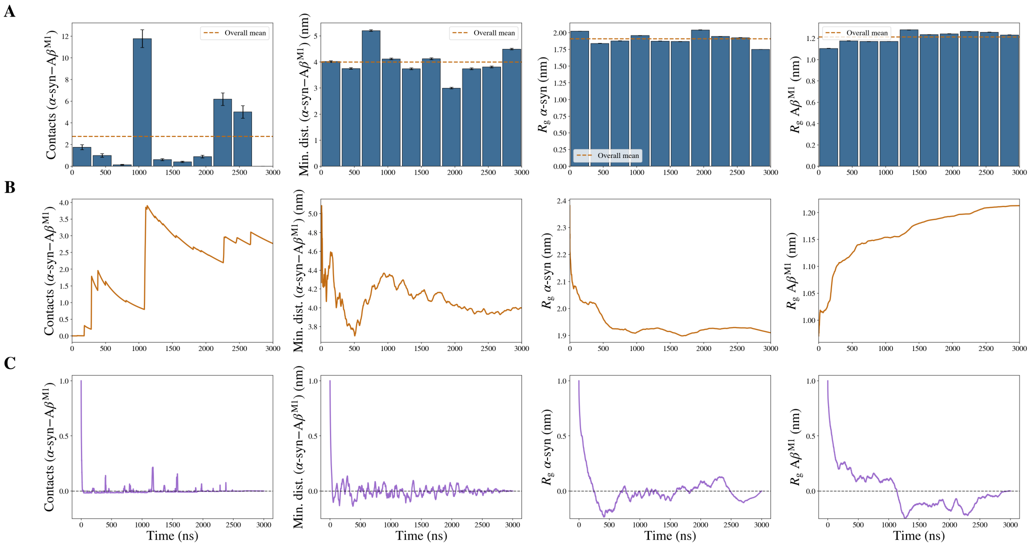


**(e)**

**Figure S2.** Sampling diagnostics for **System 2** ($\alpha$-syn + A$\beta_{42}^{\mathrm{M1}}$ and A$\beta_{42}^{\mathrm{M2}}$), replicates 1–5 (panels a–e). The $\alpha$-syn/A$\beta_{42}$-monomer interaction shown is $\alpha$-syn–A$\beta_{42}^{\mathrm{M1}}$. Within each panel, columns show (left to right) the inter-chain heavy-atom contacts, minimum inter-chain distance, $\alpha$-syn radius of gyration, and A$\beta_{42}^{\mathrm{M1}}$ radius of gyration. Rows show **(A)** block averages over ten equal blocks (error bars: standard error of the mean within each block; dashed line: overall trajectory mean), **(B)** the cumulative running average, and **(C)** the temporal autocorrelation function.

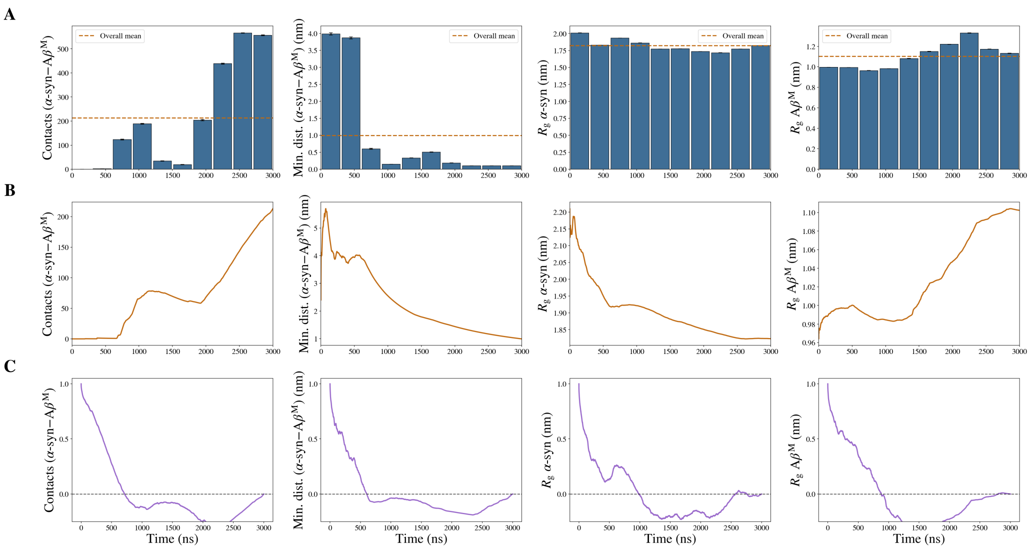
A
B
C
Overall mean
Contacts (α-syn–Aβ^M)
Min. dist. (α-syn–Aβ^M) (nm)
R_g α-syn (nm)
R_g Aβ^M (nm)
Time (ns)

(a)

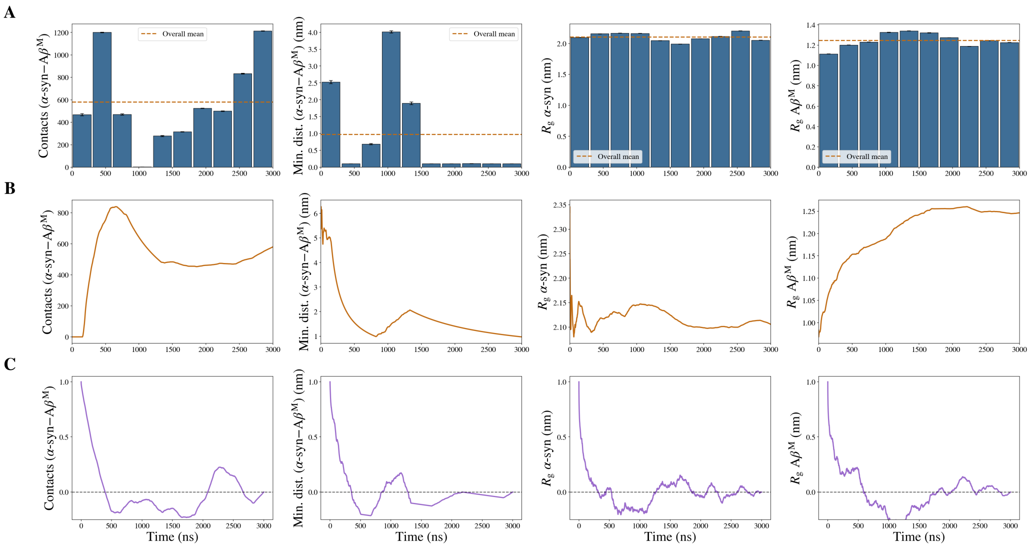
A
B
C
Overall mean
Contacts (α-syn–Aβ^M)
Min. dist. (α-syn–Aβ^M) (nm)
R_g α-syn (nm)
R_g Aβ^M (nm)
Time (ns)

(b)

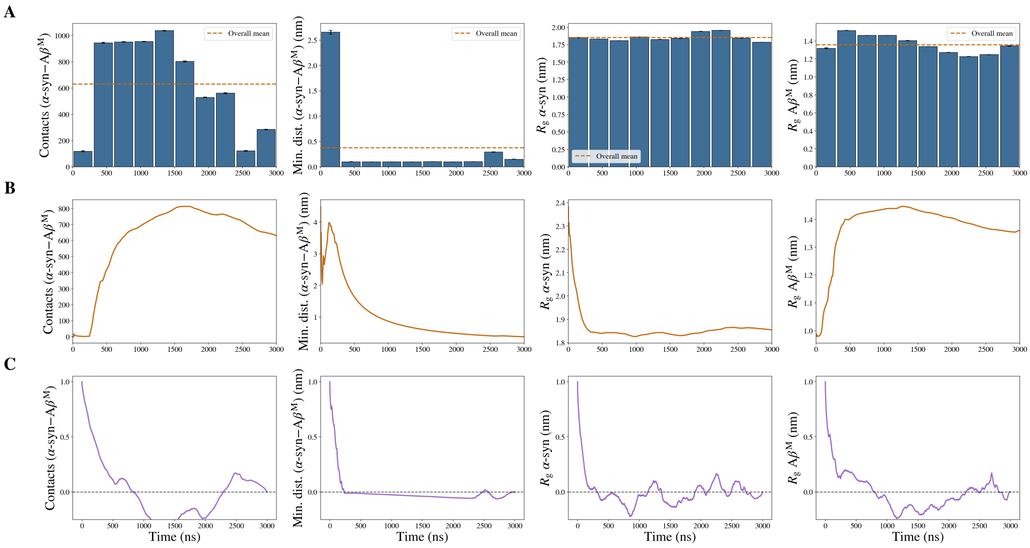
A
B
C
Overall mean
Contacts (α-syn–Aβ^M)
Min. dist. (α-syn–Aβ^M) (nm)
R_g α-syn (nm)
R_g Aβ^M (nm)
Time (ns)

(c)

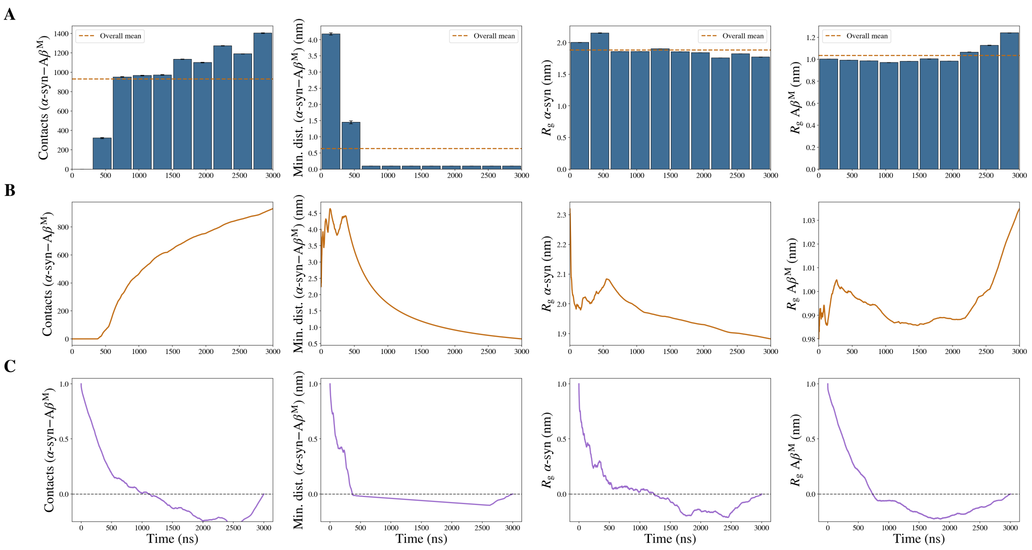


**(d)**

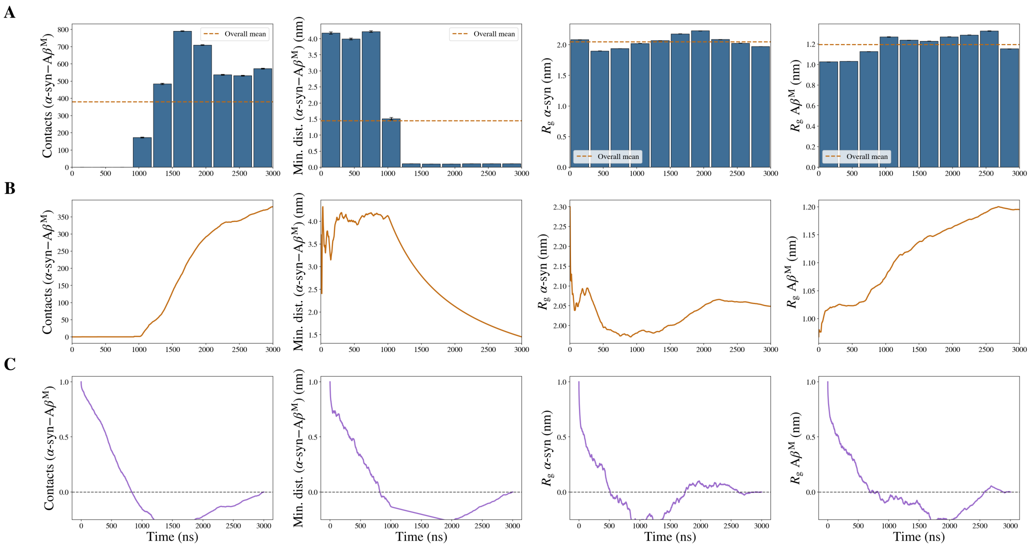


**(e)**

**Figure S3.** Sampling diagnostics for **System 3** ($\alpha$-syn + A$\beta_{42}^{\mathrm{M}}$ + A$\beta_{42}^{\mathrm{D}}$), replicates 1–5 (panels a–e). The $\alpha$-syn/A$\beta_{42}$-monomer interaction shown is $\alpha$-syn–A$\beta_{42}^{\mathrm{M}}$ (the free monomer). Within each panel, columns show (left to right) the inter-chain heavy-atom contacts, minimum inter-chain distance, $\alpha$-syn radius of gyration, and A$\beta_{42}^{\mathrm{M}}$ radius of gyration. Rows show **(A)** block averages over ten equal blocks (error bars: standard error of the mean within each block; dashed line: overall trajectory mean), **(B)** the cumulative running average, and **(C)** the temporal autocorrelation function.

### II Individual Replicate Contact Maps

This section provides detailed contact maps for individual replicates corresponding to the aggregation behaviors discussed in the main text. The contact matrices display the percentage of contact time between the specified polypeptide regions. Specifically, an active contact was defined between any two un-hydrogenated carbon atoms (@C=) within 6.0 Å. The cumulative sum of these pairwise contact events was divided by the total number of evaluated simulation frames and scaled by 100 to yield a percentage. Each figure shows the region-level contact map for all three simulation systems within a single replicate.

To quantify domain-level interactions, residue-level contact matrices were mathematically aggregated over established polypeptide structural regions based on known functional motifs [1, 2]. For A$\beta_{42}$, residues are reported using their canonical A$\beta_{42}$ numbering. The N-terminal cysteine tag is assigned position 0 and is grouped into the metal-binding domain, so that all native A$\beta_{42}$ residues retain their canonical numbers. The functional regions were defined as: Metal-binding (the N-terminal cysteine plus residues 1–16), CHC (central hydrophobic core, residues 17–21), Polar (residues 22–29), and C-term (C-terminus, residues 30–42). For $\alpha$-syn, the regions were defined as: N-term (N-terminus, residues 1–60), NAC (non-amyloid component, residues 61–95), and C-term (C-terminus, residues 96–140).

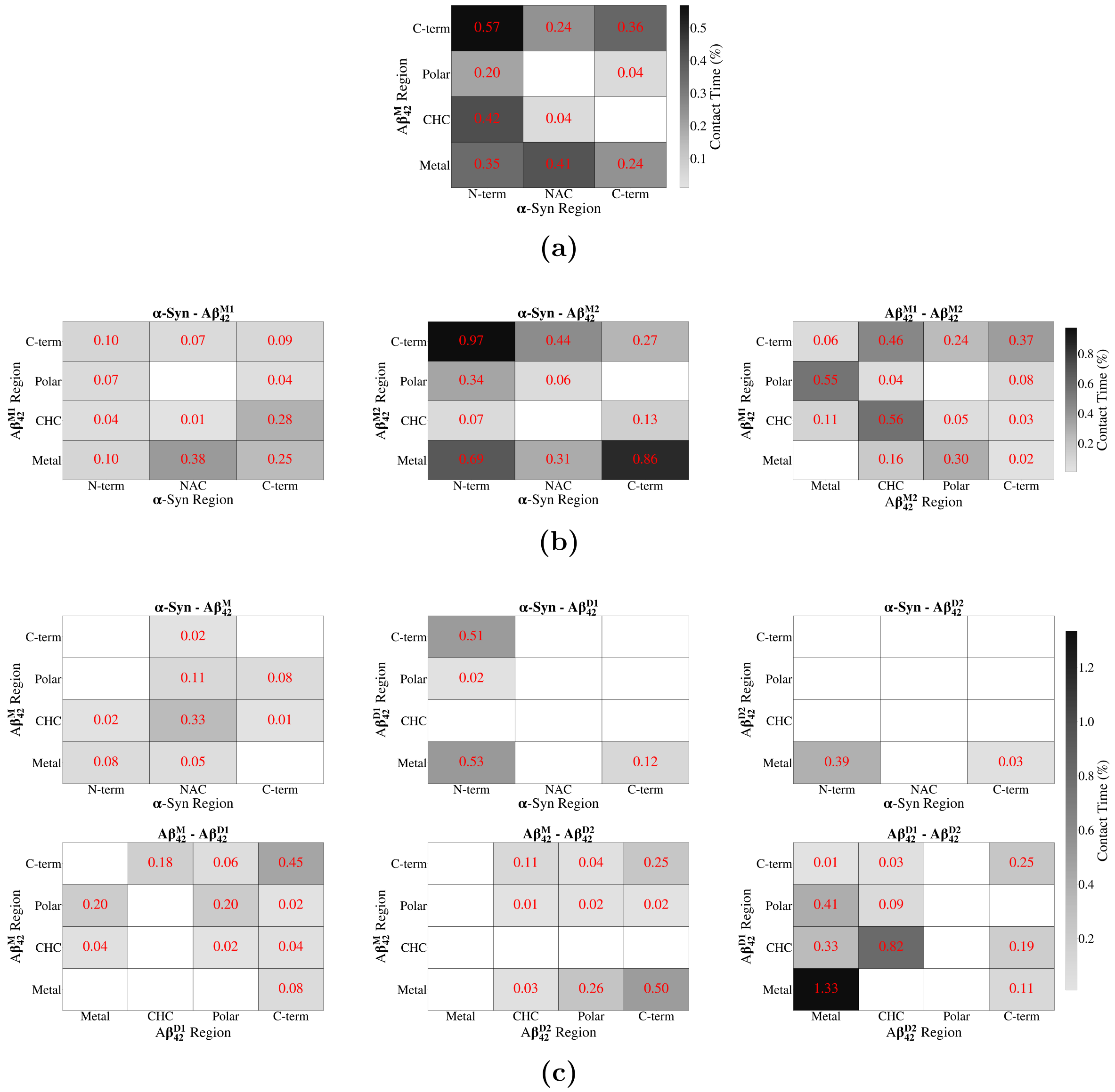


**Figure S4.** Replicate 1 contact maps across simulation systems. **(a)** System 1: $\alpha$-syn + $A\beta_{42}^{M}$. **(b)** System 2: $\alpha$-syn + $A\beta_{42}^{M1}$ and $A\beta_{42}^{M2}$. **(c)** System 3: $\alpha$-syn + $A\beta_{42}^{M}$ + $A\beta_{42}^{D}$ (D1 and D2 are labels to distinguish between the two $A\beta_{42}$ polypeptide chains within the dimer). Cell values represent the percentage of contact time between the specified regions.

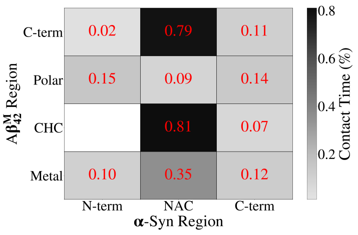


(a)

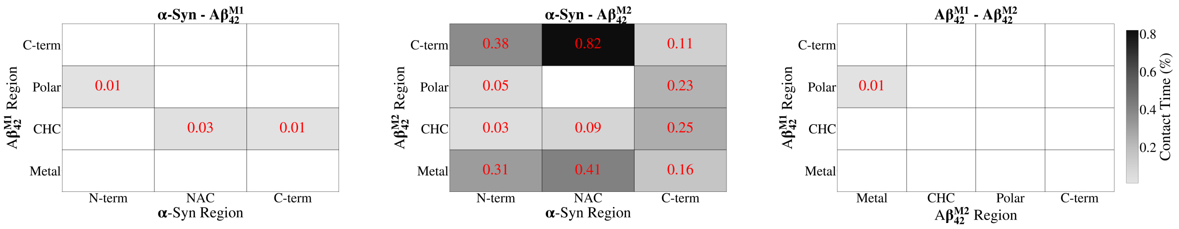


(b)

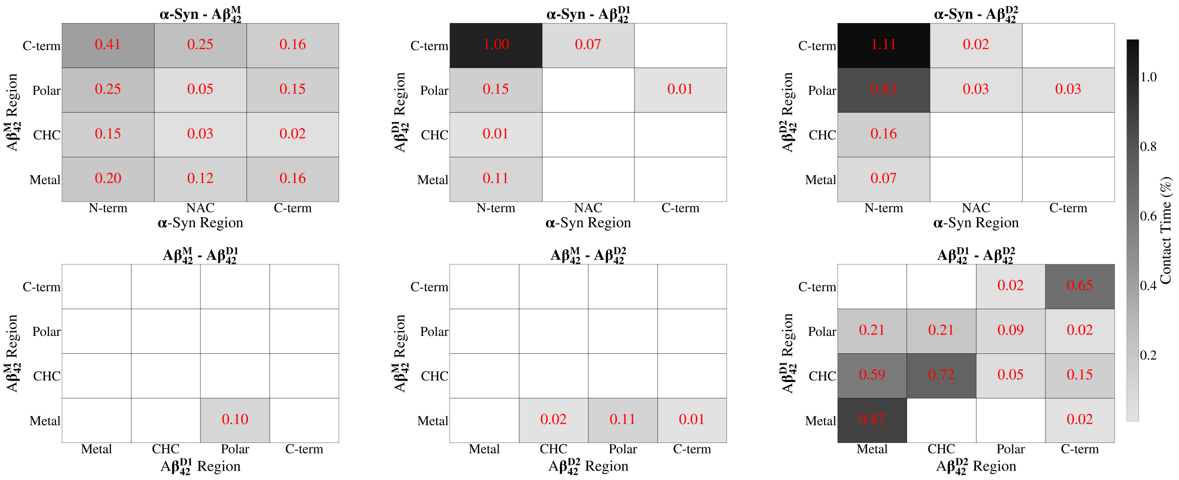


(c)

**Figure S5.** Replicate 2 contact maps across simulation systems. **(a)** System 1: $\alpha$-syn + $A\beta_{42}^{M}$. **(b)** System 2: $\alpha$-syn + $A\beta_{42}^{M1}$ and $A\beta_{42}^{M2}$. **(c)** System 3: $\alpha$-syn + $A\beta_{42}^{M}$ + $A\beta_{42}^{D}$ (D1 and D2 are labels to distinguish between the two $A\beta_{42}$ polypeptide chains within the dimer). Cell values represent the percentage of contact time between the specified regions.

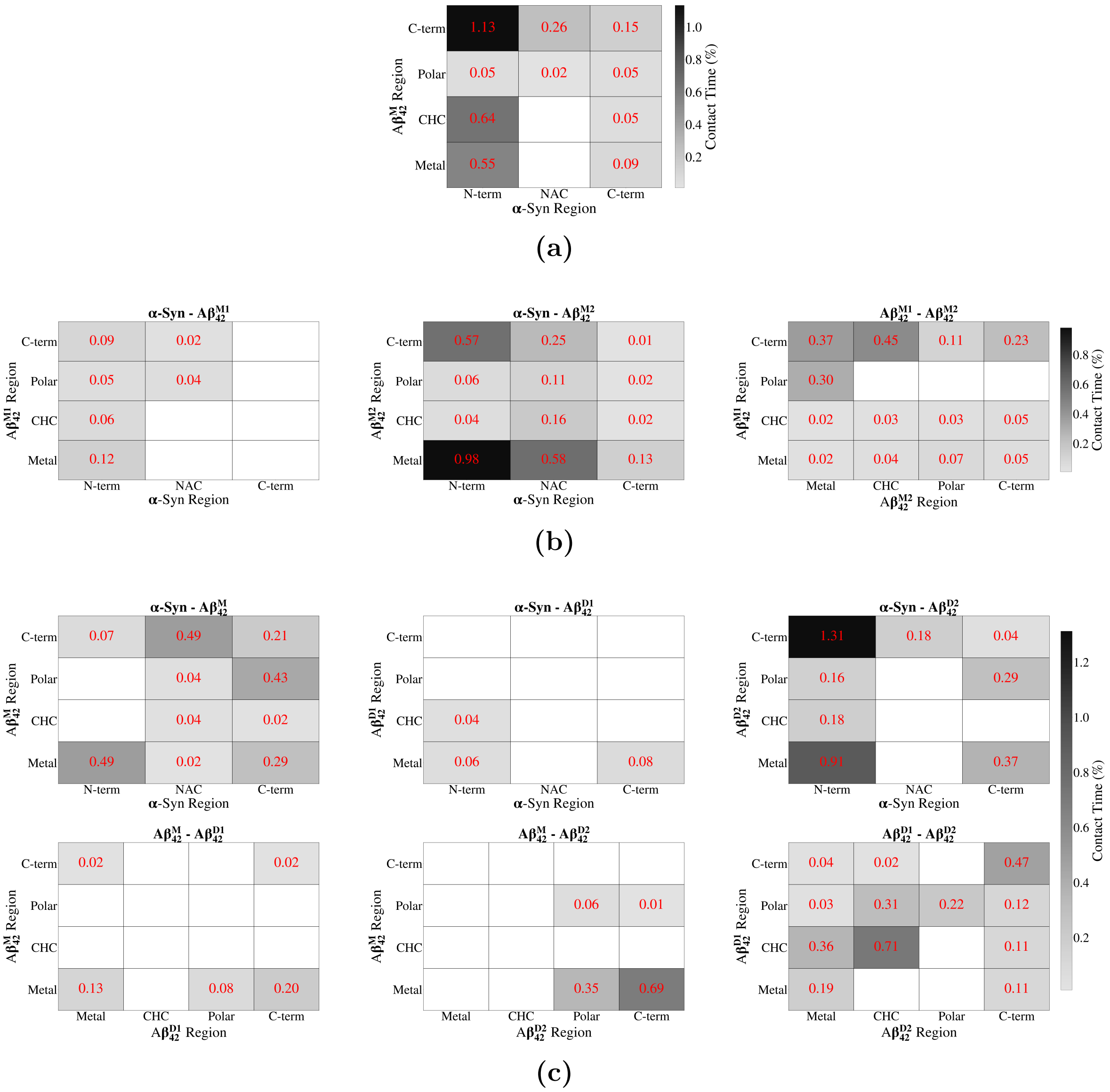


**Figure S6.** Replicate 3 contact maps across simulation systems. **(a)** System 1: $\alpha$-syn + $A\beta_{42}^{M}$. **(b)** System 2: $\alpha$-syn + $A\beta_{42}^{M1}$ and $A\beta_{42}^{M2}$. **(c)** System 3: $\alpha$-syn + $A\beta_{42}^{M}$ + $A\beta_{42}^{D}$ (D1 and D2 are labels to distinguish between the two $A\beta_{42}$ polypeptide chains within the dimer). Cell values represent the percentage of contact time between the specified regions.

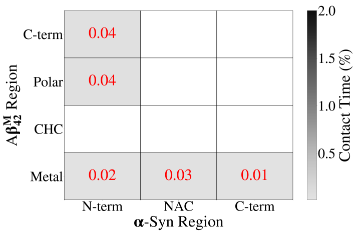


(a)

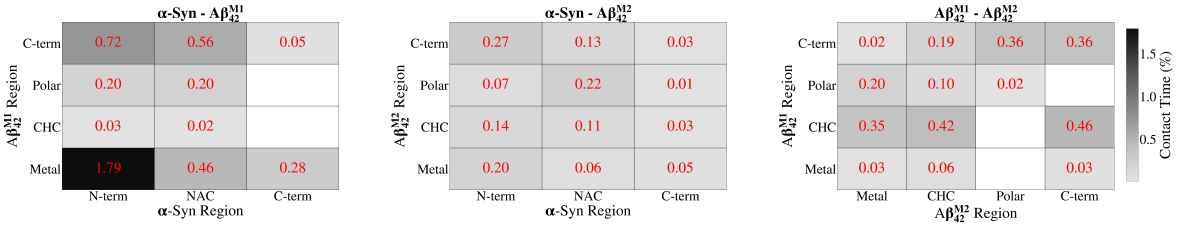


(b)

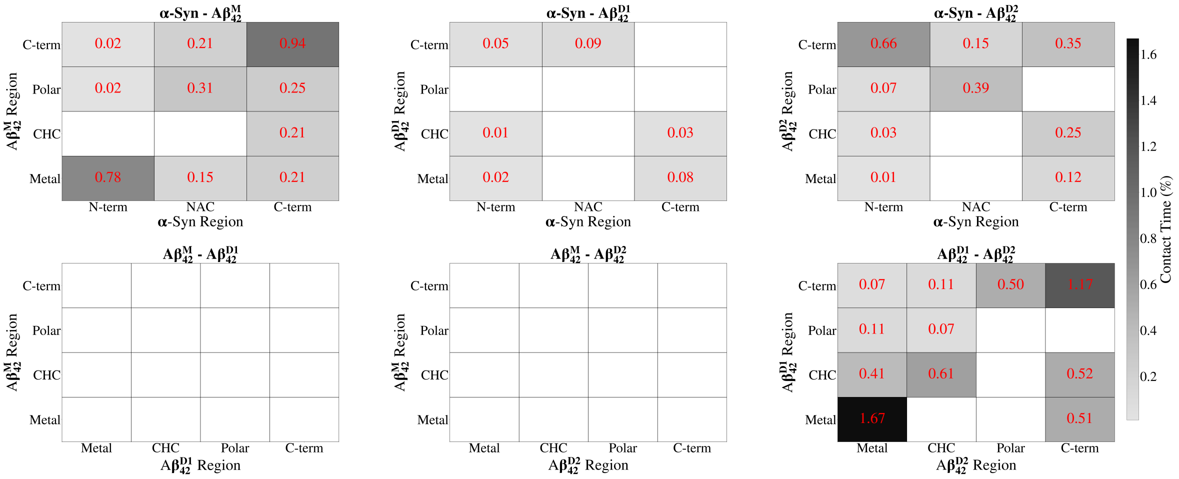


(c)

**Figure S7.** Replicate 4 contact maps across simulation systems. **(a)** System 1: $\alpha$-syn + A$\beta_{42}^{\mathrm{M}}$. **(b)** System 2: $\alpha$-syn + A$\beta_{42}^{\mathrm{M1}}$ and A$\beta_{42}^{\mathrm{M2}}$. **(c)** System 3: $\alpha$-syn + A$\beta_{42}^{\mathrm{M}}$ + A$\beta_{42}^{\mathrm{D}}$ (D1 and D2 are labels to distinguish between the two A$\beta_{42}$ polypeptide chains within the dimer). Cell values represent the percentage of contact time between the specified regions.

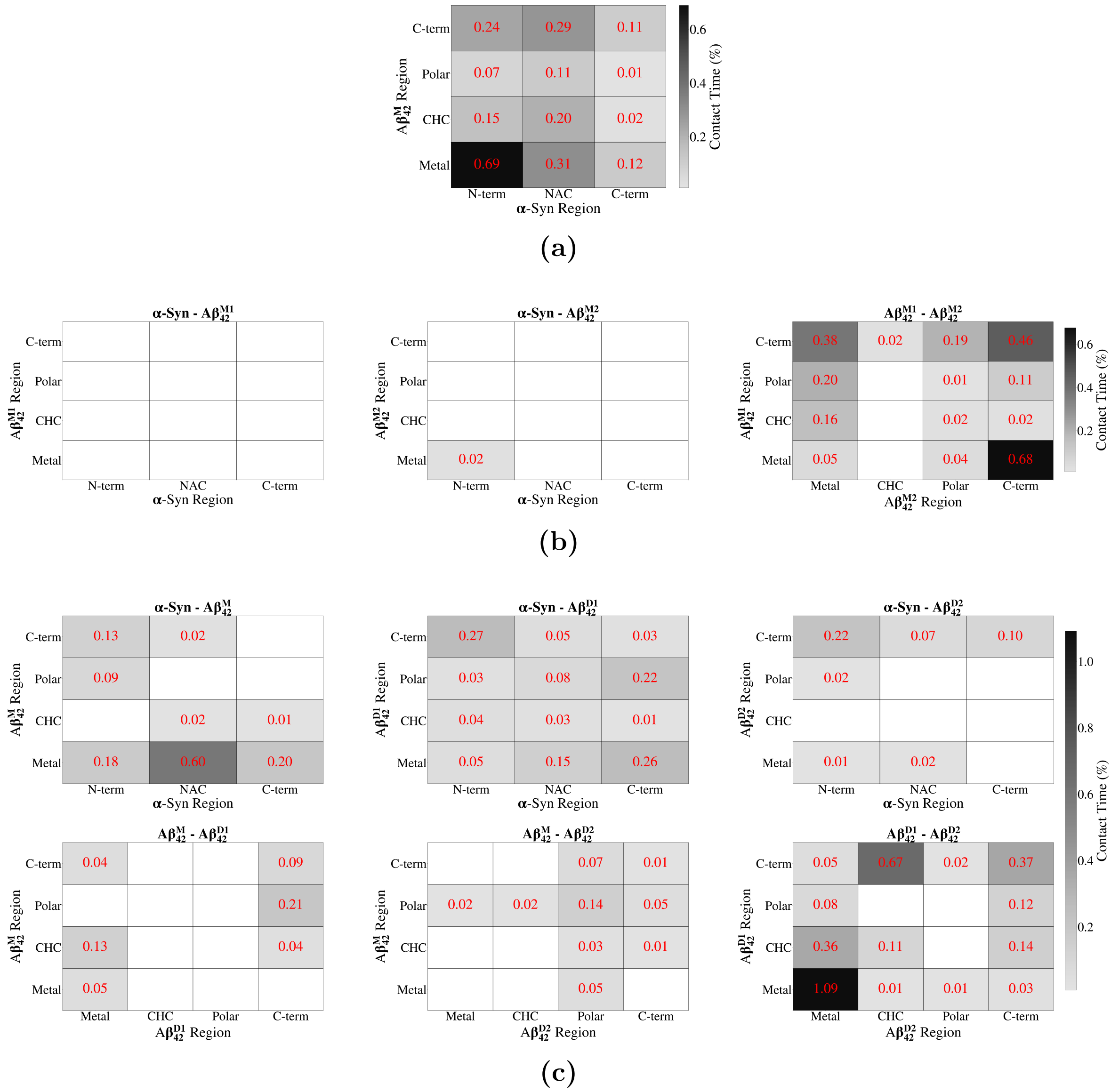


**Figure S8.** Replicate 5 contact maps across simulation systems. **(a)** System 1: $\alpha$-syn + A$\beta_{42}^{\mathrm{M}}$. **(b)** System 2: $\alpha$-syn + A$\beta_{42}^{\mathrm{M1}}$ and A$\beta_{42}^{\mathrm{M2}}$. **(c)** System 3: $\alpha$-syn + A$\beta_{42}^{\mathrm{M}}$ + A$\beta_{42}^{\mathrm{D}}$ (D1 and D2 are labels to distinguish between the two A$\beta_{42}$ polypeptide chains within the dimer). Cell values represent the percentage of contact time between the specified regions.

#### II.1 Temporal Coordination Between Contact Regions

The contact heatmaps report the time-averaged occupancy of each region-level pair but cannot indicate whether two pairs engage at the same moments in time. Figure S9 resolves this by the partial-correlation analysis described in the main document: for every pair of region-level cells, the covariation of their per-frame contact counts is quantified while controlling for the overall degree of interfacial engagement, so that a positive coefficient ($r > 0$) marks region pairs engaged concurrently and a negative coefficient ($r < 0$) marks pairs engaged alternatively. The accompanying $p_{\text{joint}}$ gives the fraction of engaged frames in which both region pairs are simultaneously in contact.

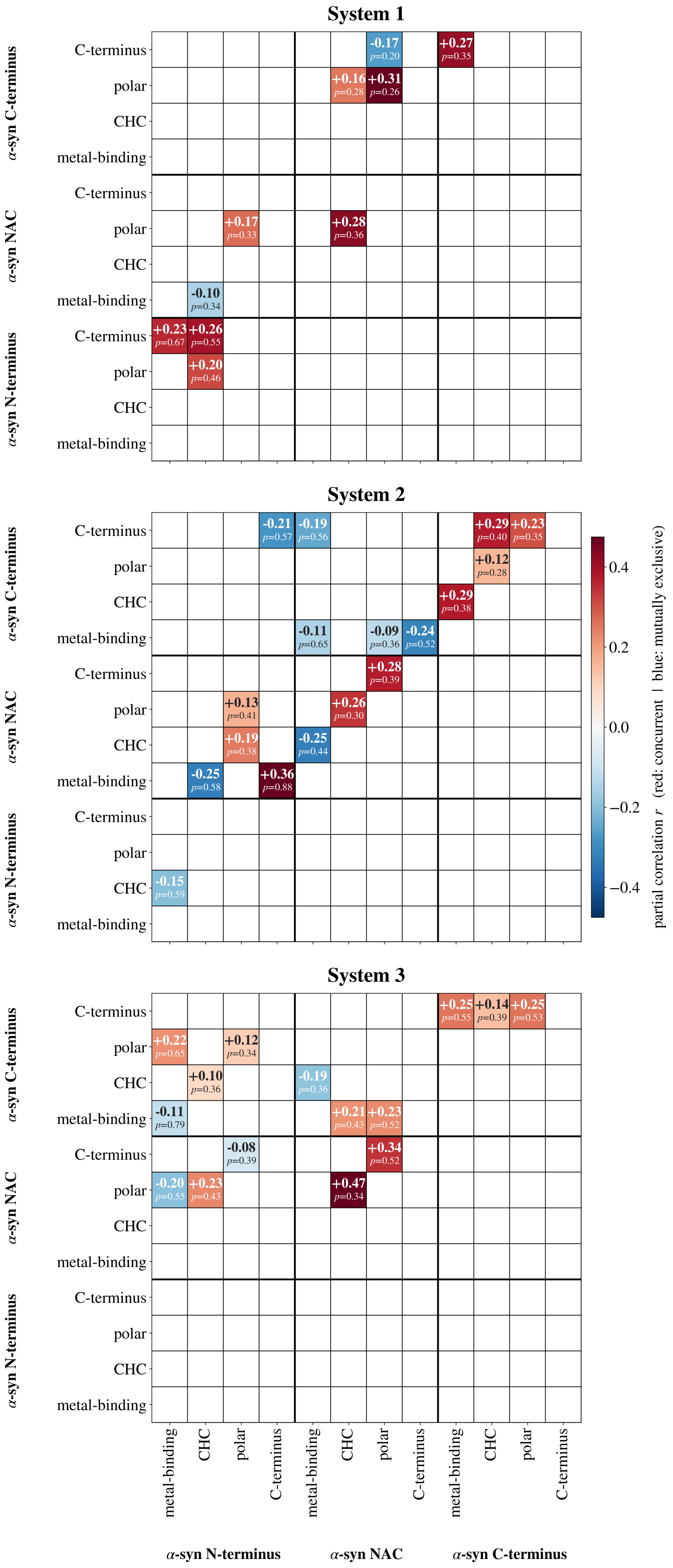


**Figure S9.** Temporal coordination between contact regions. Upper value: mean partial correlation $r$ (red, concurrent; blue, mutually exclusive); lower value: joint occupancy $p_{\text{joint}}$.

### III $A\beta_{42}$ influence on $\alpha$-syn structure

Figure S10 presents the structural ensemble distributions of $\alpha$-syn as a function of $A\beta_{42}$ contact state, complementing the structural analysis of $A\beta_{42}$ reported in the main text (Section 3.3). Across all three systems, $A\beta_{42}$ contact has a limited effect on the global $\alpha$-syn structural metrics, consistent with the observation that $\alpha$-syn acts as the dominant conformational modulator in the heterodimer rather than the modulated species.

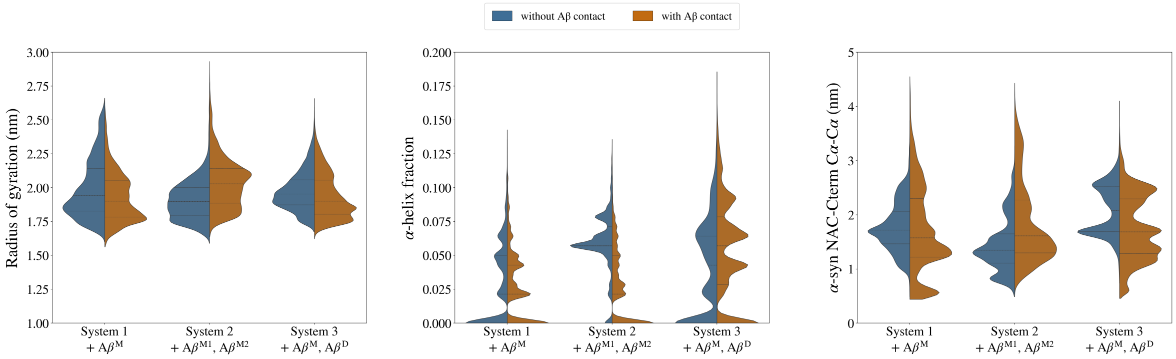


**Figure S10.** Structural ensemble distributions of $\alpha$-syn as a function of $A\beta_{42}$ contact state, across all three simulated systems. Split violins show the per-frame distribution of three structural metrics — radius of gyration ($R_g$), $\alpha$-helix fraction, and NAC-to-C-terminus C$\alpha$–C$\alpha$ distance (residues 78 and 118) — partitioned by the absence (blue) or presence (orange) of $A\beta_{42}$ contact, as defined by the dual-criteria aggregation metric. Each violin pools frames across five independent 3 µs replicates. Horizontal lines within each violin indicate the 25th percentile, median, and 75th percentile.

### IV $\beta$-Hairpin Topology Analysis

Table S1: $\beta$-Hairpin topology summary across all systems and replicates in the aqueous environment. Frames % = fraction of trajectory frames containing at least one validated target hairpin. Topology % = fraction of validated hairpin events falling in the dominant topology category, indicating how converged the family of formed hairpins is into the dominant topology. Dominant hairpin = the single most frequent strand-1/turn/strand-2 geometry within the dominant topology, expressed as residue ranges in canonical $A\beta_{42}$ numbering (N-terminal cysteine tag = position 0). Hairpin % = fraction of trajectory frames in which that specific hairpin geometry is present, indicating how committed the chain is to that exact register within the dominant topology. Em-dashes (—) indicate replicates with no validated target hairpins. Highlighted rows (System 2 $A\beta_{42}^{M1}$ replicate 5 and System 3 $A\beta_{42}^{D1}$ replicate 4) are referenced in the main text.

| Sys. | Rep | Chain | Frames (%) | Dominant topology | Topo. (%) | Dominant hairpin (s1 / turn / s2) | Hairpin (%) |
|---|---|---|---|---|---|---|---|
| | 1 | $A\beta_{42}^{M}$ | 0.8 | C-term↔C-term | 100 | 31–33 / 34–38 / 39–42 | 0.6 |
| | 2 | $A\beta_{42}^{M}$ | <0.1 | CHC↔CHC | 100 | 16–18 / 19–20 / 21–25 | <0.1 |
| 1 | 3 | $A\beta_{42}^{M}$ | 0.7 | CHC↔C-term | 100 | 21–24 / 25–27 / 28–31 | 0.5 |
| | 4 | $A\beta_{42}^{M}$ | 7.8 | CHC↔C-term | 100 | 20–23 / 24–27 / 28–32 | 5.6 |
| | 5 | $A\beta_{42}^{M}$ | 0.9 | C-term↔C-term | 99 | 35–36 / 37–39 / 40–41 | 0.4 |
| | 1 | $A\beta_{42}^{M1}$ | — | — | — | — | — |
| | 1 | $A\beta_{42}^{M2}$ | 0.1 | C-term↔C-term | 100 | 32–34 / 35–36 / 37–38 | 0.1 |
| | 2 | $A\beta_{42}^{M1}$ | 11.8 | C-term↔C-term | 100 | 32–33 / 34–35 / 36–37 | 4.0 |
| | 2 | $A\beta_{42}^{M2}$ | 9.6 | CHC↔C-term | 100 | 22–25 / 26–29 / 30–32 | 9.3 |
| | 3 | $A\beta_{42}^{M1}$ | 5.7 | C-term↔C-term | 100 | 31–32 / 33–35 / 36–37 | 4.3 |
| 2 | 3 | $A\beta_{42}^{M2}$ | — | — | — | — | — |
| | 4 | $A\beta_{42}^{M1}$ | — | — | — | — | — |
| | 4 | $A\beta_{42}^{M2}$ | — | — | — | — | — |
| | **5** | **$A\beta_{42}^{M1}$** | **76.8** | **C-term↔C-term** | **100** | **32–33 / 34–35 / 36–37** | **73.3** |
| | 5 | $A\beta_{42}^{M2}$ | — | — | — | — | — |
| | 1 | $A\beta_{42}^{M}$ | — | — | — | — | — |
| | 1 | $A\beta_{42}^{D1}$ | 42.5 | C-term↔C-term | 63 | 36–37 / 38–39 / 40–41 | 29.1 |
| | 1 | $A\beta_{42}^{D2}$ | <0.1 | C-term↔C-term | 100 | 29–32 / 33–34 / 35–41 | <0.1 |
| | 2 | $A\beta_{42}^{M}$ | 0.6 | C-term↔C-term | 88 | 36–37 / 38–39 / 40–41 | 0.4 |
| | 2 | $A\beta_{42}^{D1}$ | 95.6 | C-term↔C-term | 100 | 35–37 / 38–39 / 40–42 | 64.3 |
| | 2 | $A\beta_{42}^{D2}$ | — | — | — | — | — |
| | 3 | $A\beta_{42}^{M}$ | 3.0 | C-term↔C-term | 100 | 31–34 / 35–37 / 38–42 | 1.7 |
| 3 | 3 | $A\beta_{42}^{D1}$ | 32.8 | C-term↔C-term | 99 | 36–37 / 38–39 / 40–41 | 26.8 |
| | 3 | $A\beta_{42}^{D2}$ | — | — | — | — | — |
| | 4 | $A\beta_{42}^{M}$ | 1.5 | CHC↔CHC | 100 | 16–18 / 19–20 / 21–25 | 1.5 |
| | **4** | **$A\beta_{42}^{D1}$** | **25.4** | **CHC↔C-term** | **65** | **20–24 / 25–27 / 28–32** | **16.0** |
| | 4 | $A\beta_{42}^{D2}$ | — | — | — | — | — |
| | 5 | $A\beta_{42}^{M}$ | <0.1 | C-term↔C-term | 100 | 32–34 / 35–36 / 37–38 | <0.1 |
| | 5 | $A\beta_{42}^{D1}$ | 93.4 | C-term↔C-term | 91 | 32–37 / 38–39 / 40–42 | 59.3 |
| | 5 | $A\beta_{42}^{D2}$ | 8.1 | CHC↔C-term | 69 | 34–36 / 37–39 / 40–42 | 2.0 |

### V Time-Resolved Configurational Entropy

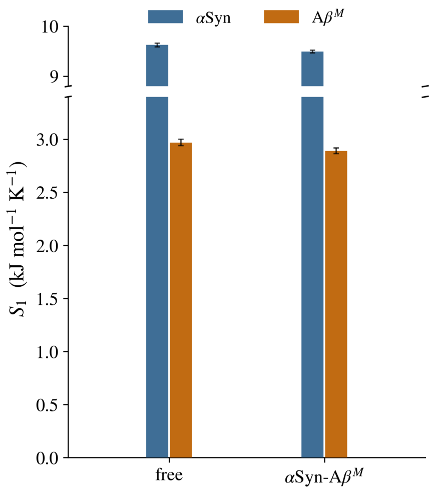


**(a)** System 1

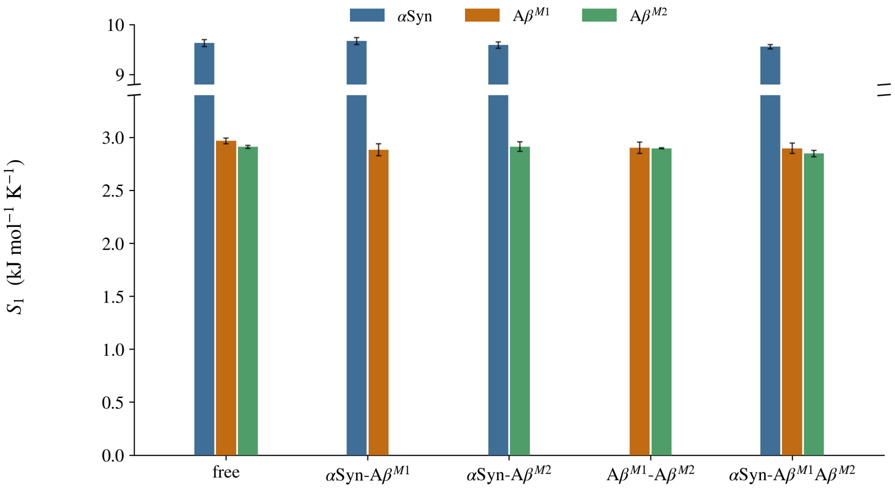


**(b)** System 2

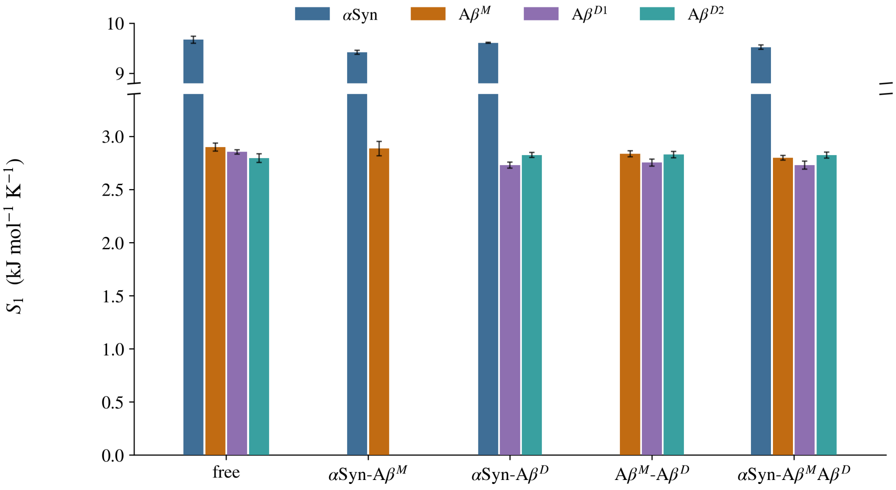


**(c)** System 3

**Figure S12.** First-order backbone conformational entropy ($S_1$) profiles resolved across distinct aggregation states in aqueous solution. Bar summaries depict the mean of per-replicate median windowed entropies (± SEM across 5 independent trajectories) computed using a 20 ns moving sampling window.
